# Structure of Infective Getah Virus at 2.8 Å-resolution Determined by Cryo-EM

**DOI:** 10.1101/2021.07.16.452580

**Authors:** Aojie Wang, Feng Zhou, Congcong Liu, Dongsheng Gao, Ruxi Qi, Yiheng Yin, Sheng Liu, Yuanzhu Gao, Lutang Fu, Yinhe Xia, Yawei Xu, Chuanqing Wang, Zheng Liu

**Affiliations:** Cryo-electron Microscopy Center, Southern University of Science and Technology, Shenzhen, Guangdong, China; College of Animal Science and Veterinary Medicine, Henan Agricultural University, Zhengzhou, Henan, China; Institute for Hepatology, National Clinical Research Center for Infectious Disease, Shenzhen Third People’s Hospital, Shenzhen, Guangdong, China; Department of Cardiology, Shanghai Tenth People’s Hospital, and Pan-Vascular Research Institute, Heart, Lung, and Blood Center, Tongji University School of Medicine, Shanghai, China

## Abstract

Getah virus (GETV), a member of genus *alphavirus*, is a mosquito-borne pathogen that can cause pyrexia and reproductive losses in animals. Although antibodies to GETV have been found in over 10% of healthy people, there are no reports of clinical symptom associated with GETV. The biological and pathological properties of GETV are largely unknown. Here, we present the structure of infective GETV at a resolution of 2.8 Å with the capsid protein and the envelope glycoproteins E1 and E2. We have identified numerous glycosylation and S-acylation sites in E1 and E2. The surface-exposed glycans revealed their impact on the viral immune evasion and host cell invasion. The S-acylation sites involve in stabilizing the transmembrane assembly. In addition, a cholesterol and a phospholipid molecule are observed in a transmembrane hydrophobic pocket, together with two more cholesterols surrounding the pocket. The structural information will assist structure-based antiviral and vaccine design.

## Introduction

Getah virus (GETV) is a mosquito-borne arbovirus and belongs Semliki Forest group of the *Alphavirus* genus within the *Togaviridae* family (Lvov et al., 2015). Alongside GETV, members in Semliki Forest Group include Chikungunya virus (CHIKV), Semliki Forest virus (SFV), Mayaro virus (MAYV), Una virus (UNAV), Bebaru virus (BEBV), and O’nyong-nyong virus (ONNV). Among them, CHIKV, ONNV, and MAYV have been reported to cause severe and mortally dangerous infection diseases in human (Hua and Combe, 2017; Rezza et al., 2017; Tesh et al., 1999).

GETV was first isolated in Malaysia in 1955 from *Culex gelidus* mosquitoes (Berge, 1975), and was found to have wide distribution in worldwide. The primary hosts of GETV include pigs, cattle, foxes and horses. The first outbreak of GETV among racehorses occurred in Japan in 1978 and outbreaks have re-emerged several times in Japan (Bannai et al., 2015). The infected horses exhibited pyrexia, urticarial rash on various portions of the body, and edema of the hind legs (Kamada et al., 1980). The emergence of GETV in China was found in 2017 in swine farms, resulting in the death of the newborn piglets 5-10 days after birth and in pregnant sows having stillbirths or foetal mummies (Yang et al., 2018). GETV was also reported to infect beef cattle (Liu et al., 2019), blue fox (Shi et al., 2019), and wild boars (Sugiyama et al., 2009). Neutralizing antibodies of GETV have been detected in a variety of vertebrate species, suggesting that more domestic animals could act as reservoir hosts (Li et al., 2019). Although the pathogenicity of GETV in humans has not yet been identified, seroepidemiologic investigations have shown that some individuals with a febrile illness have significantly higher GETV antibody titers than in healthy people (Li et al., 1992), suggesting an association of GETV with human diseases.

Like a typical alphavirus, GETV is a lipid-enveloped, positive-sense single-stranded RNA virus (Lvov et al., 2015). Mature virions of alphaviruses are spherical particles with a diameter of ∼70 nm. The 11-kb genome of GETV encodes two polyproteins. Among them, one polyprotein consists of four non-structural proteins (nsP1-nsP4), and another one polyprotein composes five structural proteins, in order from the N-terminal to the C-terminal, capsid, E3, E2, 6K and E1 (Rayner et al., 2002). The structures of alphaviruses have been well studied by using cryo-electron microscopy (cryo-EM). These include the structures of Barmah forest virus (BFV), EEEV, WEEV, VEEV, CHIKV, SINV, and MAYV, ranging from 3.5-13 Å in resolution; there is also structures of CHIKV with its receptor MXRA8 (Basore et al., 2019; Chen et al., 2020; Chen et al., 2018; Hasan et al., 2018; Kostyuchenko et al., 2011; Ribeiro-Filho et al., 2021; Sherman and Weaver, 2010; Yap et al., 2017; Zhang et al., 2011). These structures reveal a typical architecture for alphavirus organization: viralRNA occupies the center of the particle and extends to approximately 140-Å radially, and the RNA is surrounded by the capsid protein shell ranging between 140 to 200-Å radially; the lipid membrane shell (200 to 255-Å radially) separating the capsid and the outer glycoprotein shell, the outer shell and spikes protruding outward (255 to 350-Å radially) are formed by the E1 and E2 glycoproteins. In addition, high-resolution maps have assisted the building of atomic models for the capsid, glycoproteins E1 and E2 (Chen et al., 2018; Hasan et al., 2018; Ribeiro-Filho et al., 2021; Zhang et al., 2011). Moreover, Mxra8, the receptor of CHIKV, was found to bind into a cleft created by two adjacent CHIKV E1-E2 heterodimers (Basore et al., 2019).

In a previous study, we have isolated a new GETV strain, V1 strain, from pregnant sows had abortions, and have demonstrated the V1 strain of GETV had a strong cytopathic effect and higher proliferation titer of vaccine (Zhou et al., 2020). In this study, we utilized mice model to investigate the infectiousness and pathogenicity of mature GETV. To gain structural insight into GETV, we determined the structure of the mature GETV V1 virion at 2.8 Å-resolution by cryo-EM, the highest resolution for an alphavirus.

## Results

### GETV is a mosquito-borne arbovirus

GETV appears to be maintained in a natural cycle between mosquitoes and various vertebrate hosts (Fukunaga et al., 2000). There is also suspicion that GETV can be transmitted directly between horses, likely via aerosols or by direct contact (Kamada et al., 1980). There is no clear evidence how GETV transmits and infects by mosquitoes. Therefore, we first investigated the transmission and infectiousness of GETV using a mouse model. As shown in **Figure S1A**, mice (2-day old) were inoculated oronasally with 10μl (10^6^ TCID_50_/mL, 50% tissue culture infectious dose) of the GETV V1 strain, and were then housed in a cage with their GETV-free littermates. Total RNA was extracted from tissue, including spleen, lung, cerebral cortex, and various lymph nodes at 5 days post inoculation (DPI), and subjected to RT-PCR for GETV detection (**Figures S1G** and **S1H**). The uninfected mice remained negative for GETV (**Figure S1A**). When male adult inoculated mice (2-month old) were housed with their uninfected male littermates (**Figure S1B**), all mice tested GETV-positive at 5 DPI. Multiple wounds were found in various part of their body, indicating that the transmission of GETV occurred through scratching and biting.

We also conducted experiments based on a special device we designed to connect two cages by a pipe; we placed screens at the two ends of the pipe to separate the mice but to allow mosquitoes to freely fly between the cages (**Figure S1C**). In a control test shown in **Figure S1D**, GETV-free mice in the two cages lived with mosquitoes and tested negative after five days, indicating that the mosquitoes used in our study were GETV-free. Mice were oronasally inoculated with GETV, then placed with their non-inoculated littermates in the left-hand cage, additional GETV-free littermates were placed in the right-hand cage, and mosquitoes were introduced into the device (**Figure S1E**). At 5 DPI, all of the mice in the left-hand cage were GETV-positive, and 15% of the mice in the right-hand cage were positive as well. In another control test, no mosquitoes were introduced, all of the non-inoculated mice in the right-hand cage remained negative after five days (**Figure S1F**). These results confirmed that GETV is a mosquito-borne arbovirus, not an airborne virus. GETV mainly spread by mosquito biting, travel in the blood between mouse and mosquitos. GETV can also transmit between mouses, most likely through direct contact with blood and body fluids.

### Pathogenicity of the GETV V1 strain

GETV has been reported in reproductive losses in pregnant sows (Yang et al., 2018). In the present study, we investigated the pathogenicity of GETV in pregnant mice. Mice were inoculated oronasally with 100 μl (10^6^ TCID_50_/mL) of the GETV V1 strain in early- (embryonic day 6, E6), middle- (E10), and late-gestation (E14). The inoculated E6 group mice displayed severe abortion (**Figure S1K**) and delivered foetal mummies (**Figures S1L**). The E10 group delivered mainly mummies or stillbirths, but also delivered a small proportion of health pups (**Figures S1K**–**S1O**). The E14 mice delivered mixture of mummies, stillbirths and live infants, although some newborn mice were weak (**Figures S1L**–**S1O**). In the contrast, no birth defect was detected in the mice inoculated with DMEM at E6 (**Figures S1O** and **S1P**). Surprisingly, although GETV-infected pregnant mice displayed serious reproductive losses, no other disease symptom was discovered, both miscarriage and maternity mice apparently normal. These results suggest that when pregnant mice are infected with GETV, placental and fetal infection occur, eventually causing fetal damage, such as abortion, mummies or stillbirths.

Next, we investigated the pathogenicity of GETV in newborn mice. 3-day old mice were inoculated oronasally with 10μl (10^6^ TCID_50_/mL) of the GETV V1 strain, and clinical signs were observed daily. At 4 DPI, the inoculated mice displayed mobility impairments in pelvic limbs (**Figures 1A** and **Movie S1**); by 8 DPI, all of the inoculated mice displayed paralysis, mainly in pelvic limbs. Near 20% inoculated mice died at 6 DPI (2 days after their paralysis manifestation), and all of the inoculated mice were dead by 12 DPI (**Figure 1B**). On the opposite, no mice had paralysis nor death in the control group (**Figures 1A** and **1B**). Immunohistochemistry staining revealed that abundant GETV antigens were distributed in the hippocampal dentate gyrus of the brain and in the spinal cord under lumbar vertebrae in the inoculated newborn mice (**Figure 1C**). Interestingly, when 7-day old newborn or 2-month old adult mice were inoculated with GETV, no such clinical signs were observed. This suggests that GETV is only transmitted into brain before the blood-brain barrier is established (Oliver et al., 1997); it also suggests that the mobility impairments in pelvic limbs are possibly caused by viral infection in the nervous system.

**Figure 1.**
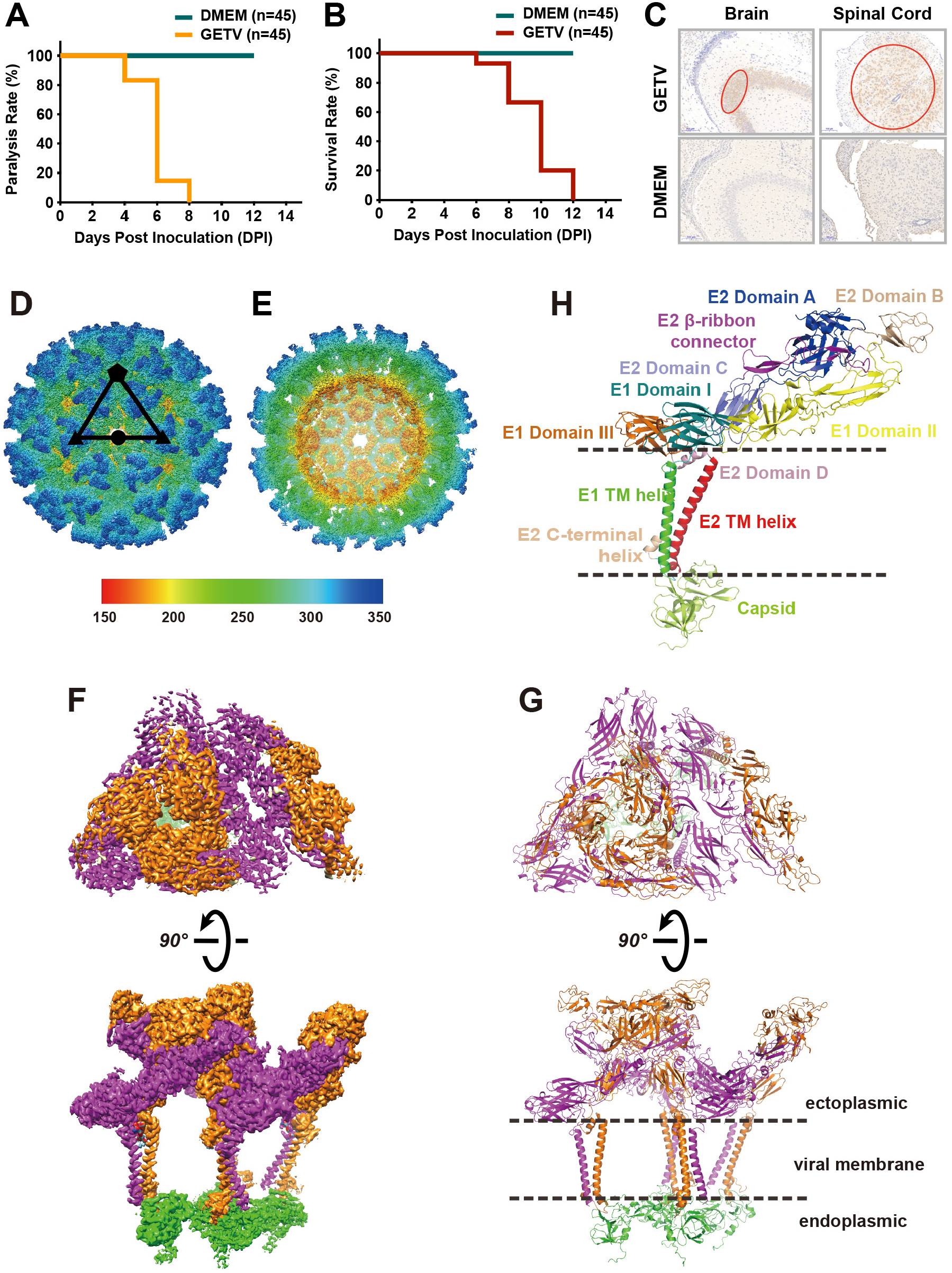
Overall structure of the infectious GETV virion determined by cryo-EM at 2.8 Å-resolution. Ninety 3-day old mice were divided into two groups, inoculated oronasally with GETV V1 strain or equal amount of DMEM. **A**, Motor disabilities displayed in GETV inoculated mice at 4 DPI; by 8 DPI, all of the inoculated mice displayed paralysis. **B**, Inoculated mice died at 6 DPI, and all of the inoculated were dead by 12 DPI. **C**, tissue from hippocampal dentate gyrus in the brain and from spinal cord under lumbar vertebrae were subjected for immune-histochemistry. There were abundant GETV antigens distributed in brain and spinal cord under lumbar vertebrae in the inoculated newborn mice (highlighted in red circles). Cryo-EM structure of GETV at 2.8 Å-resolution: **D**, GETV virion showing the external surface with assigned symmetry axes. **E**, Central cross-section of the GETV density map. **F** and **G**, Density map and atomic model of one asymmetric unit. The E1, E2, and capsid proteins are colored separately in magenta, orange and green. **H**, Atomic model of E1-E2-capsid heterotrimer. Different subdomains of E1 (Domain I, II, III, and TM helix) and E2 (Domain A, B, C, D, TM helix, and C-terminal helix) are shown, following the previous definition for alphaviruses (Lescar et al., 2001; Voss et al., 2010).

### Overall structure of GETV

To gain structural insight into GETV, we determined the structure of the mature GETV virion by cryo-EM. The whole structure of GETV was determined to 4.1 Å using Relion3.1 and JSPR software (**Figure S2**). To overcome the heterogeneity of the ∼70nm virus, a block-based reconstruction method was adopted (**Figures S2** and **S3**) (Zhu et al., 2018). Three blocks (5/3/2-fold symmetry axis) of GETV cryo-EM maps at 2.81/2.92/2.85 Å were acquired, respectively (**Figure S3** and **Table S1**). The density maps of the capsid in the pentamers are sufficient to observe the bulk residues. However, the densities for the capsid hexamers were still intermittent. The quality of this map was improved upon averaging of 15 equivalent capsid densities from the 3- and 2-fold symmetry axis blocks. After averaging, the correlation coefficient between the densities of pentamer and hexamer was 0.92 (**Figure S4**). By combining these three high-resolution densities, the over-all structure of GETV was obtained (**Figures 1D and 1E**). Similar to other alphaviruses, GETV has an icosahedral symmetry (T=4), comprising 60 quasi-three-fold symmetry trimers (Q-trimer) and 20 icosahedral three-fold symmetry trimers (I-trimer) (Chen et al., 2018). There are 240 capsid proteins connecting with the corresponding E2 proteins. Each asymmetric unit (ASU, **Figures 1F** and **1G**) consists four E1-E2-capsid heterotrimers (**Figure 1H**), with three heterotrimers forming one unabridged Q-trimer connecting to one heterotrimers from I-trimer.

The atomic model of the E1-E2-capsid heterotrimers includes the full-length sequences of E1 and E2, as well as the residues 111-268 of the capsid protein. The density for residues 1-110 of the pentamer capsid was not evidence in the cryo-EM map, suggesting a flexible architecture in the core of the virion. Notably, we subjected the sequence to the new released AlohaFold 2 (Jumper et al., 2021), only flexible loop structures were predicted.

The majority of the E1 ectodomain (Lescar et al., 2001) of GETV is divided into 3 subdomains (**Figure 1H**): domain I (residues 1-36, 130-164, and 274-281), domain II (residues 37-129 and 165-273; of which residues 86-96 forming a hydrophobic fusion loop that inserts into the canyon formed by E2’s A and B subdomains), and domain III (residues 282-379). Each of E1’s three domains have a secondary structure of comprising antiparallel β strands. The remaining ectodomain amino acids (residues 380-401) comprise the stem loop, which connects domain III and the transmembrane (TM) helix. The E2 ectodomain (Voss et al., 2010) comprises three subdomains (**Figure 1H**): domain A (residues 1-143), domain B (173-242), and domain C (270-344). In addition, E2’s residues 144-172 and 243-269 form the β-ribbon.

### Molecular Interactions in the E1-E2-capsid heterotrimer

The E1, E2, and capsid proteins form the fundamental heterotrimer (**Figure 1H**). There are five regions where the E1 and E2 proteins interact (**Figure 2A**). In the first region, E1’s V55, S57, P58, and S66 residues interact with the E2’s residues S239, R244, Q247, and S249 (**Figure 2B**). This first region is located near the tip of E1’s domain II and the E2’s β-ribbon, which is similar to the other alphaviruses (e.g., E1 S57 with E2 H164 in SINV (Chen et al., 2018); and E1 S57 with E2 H170 in CHIKV (Voss et al., 2010)). Other residues in this interaction region include E1’s A92 and Y93 and E2’s H226. These three residues are located at the interface between the E1’s fusion loop and E2 domain B, similar to reports for SFV (Roussel et al., 2006) and CHIKV (Voss et al., 2010).

**Figure 2.**
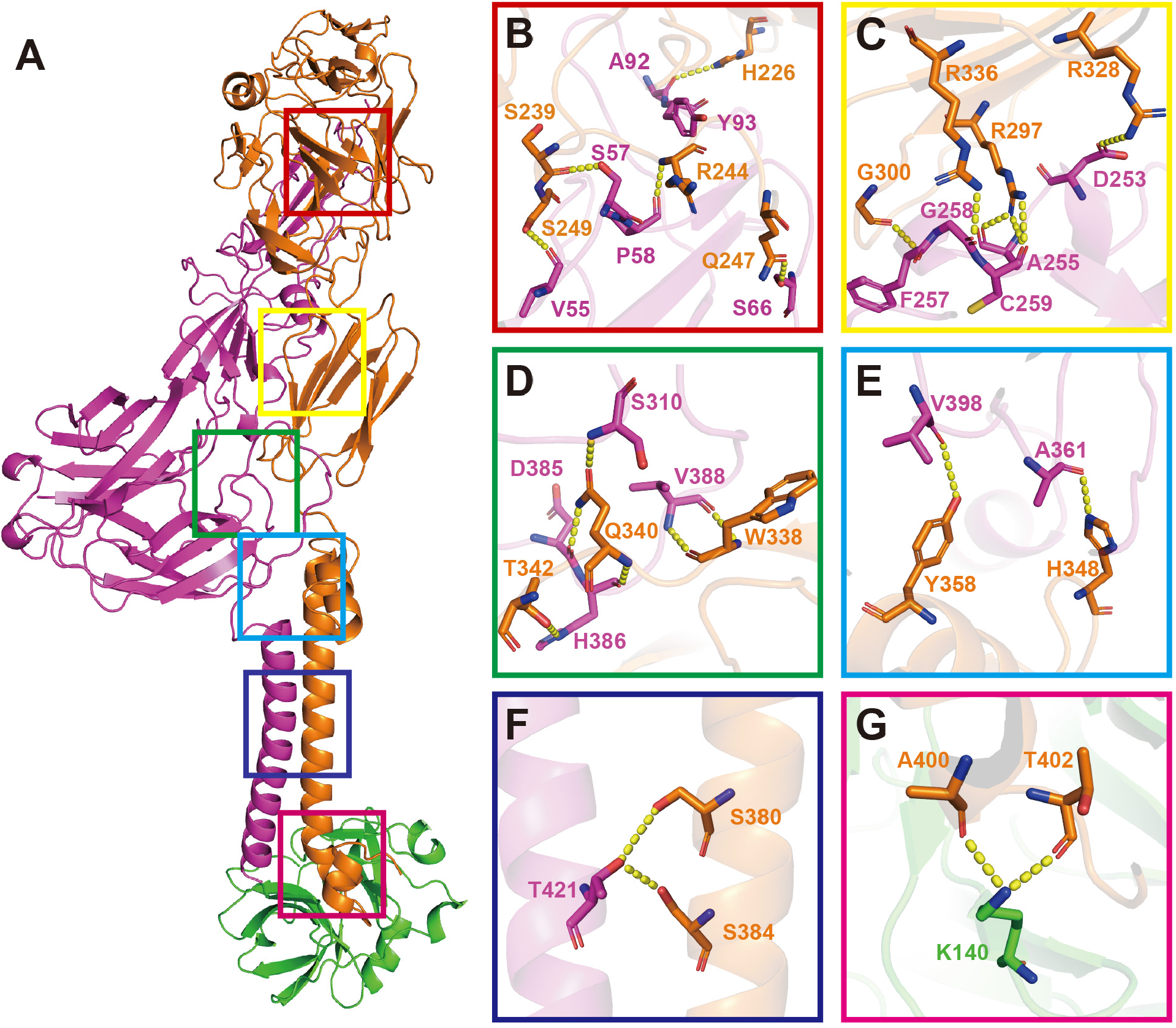
Interaction of GETV protein subdomains in the E1-E2-Capsid heterotrimer. **A**, Distribution of interaction regions between E1 and E2 and between E2 and the capsid protein. **B – F**, Zoomed-in views from A of the interaction regions between E1 and E2. **G**, Zoomed-in view from A of the interaction region between E2 and the capsid protein. The yellow dashed lines indicate the distance between the atoms involved in the interaction (the cut-off distance is 3.5 Å).

In the second interaction region, E1’s D253, A255, F257, G258, and C259 residues interact with E2’s R297, G300, R329, and R336 residues (**Figure 2C**). These interactions occur near the tail of E1’s domain II and E2’s domain C. In the third interaction region – locates at E1’s stem loop, loop of domain III and the tail loop of E2 domain C – E1’s of S310, D385, H386, and V388 residues interact with E2’s W338, Q340, and T342 residues (**Figure 2D**). Together with the second interaction region, these two areas may be required for E1/E2 dimerization. The E1 and E2 proteins pulled together through these interaction areas formed near by the stem loop of E1.

The fourth interaction region is located at E1’s stem loop, loop of domain II and E2 domain D, nearing the hydrophobic pocket (**Figure 2E**). E1’s A361 and V398 residues interact with E2’s H348 and Y358. This region is thought to be functionally important for the stability of a hydrophobic pocket (see subsection below). The fifth interaction region involves E1’s T421 and E2’s S380 and S384 (**Figure 2F**). This region is located between the E1 and E2 TM helixes and stabilizes these two helixes. There is also an interaction region between E2 and the capsid protein that is responsible for fixing the E1-E2 heterodimer on the capsid. E2’s A400, T402 residues interact with capsid protein’s K140 (**Figure 2G**). Almost all of these residues described above are conserved in the alphaviruses (**Figures S4**-**S7**).

### Intra- and Inter-ASU interactions increase virion stability

The E1-E2-capsid heterotrimer forms the Q-trimer and I-trimer. The ASU is formed by one Q-trimer and one third of I-trimer. Within the Q-trimer, there are three regions for inter-molecular interactions (**Figures 3A, 3B, 3C**, and **3F**). The E2-E2 homodimeric interaction occurs between the middle of domain A in the first E2 protein and a region of the second E2 protein comprising the joint of domains A and C, this region is positioned near by the tail of β-ribbon domain (**Figure 3B**). The second interaction locates at the helix on E1’s domain II and the E2’s domain C to stabilize the Q-trimer (**Figure 3C**). Residues K151 and D148 in one capsid interact with D120 and K122 in the neighboring capsid (**Figure 3F**). In addition, E1 of I-trimer interact with the adjacent E1 of the Q-trimer at two regions (**Figures 3D** and **3E**). E1’s T153 and E151 in domain I of the I-trimer interact with residues Y192 and K123 at the loop of E1’s domain II in the Q-trimer (**Figure 3D**). Residues T41 and N43 locate at the β sheets of domain II in two E1s symmetrically interact each other (**Figure 3E**).

**Figure 3.**
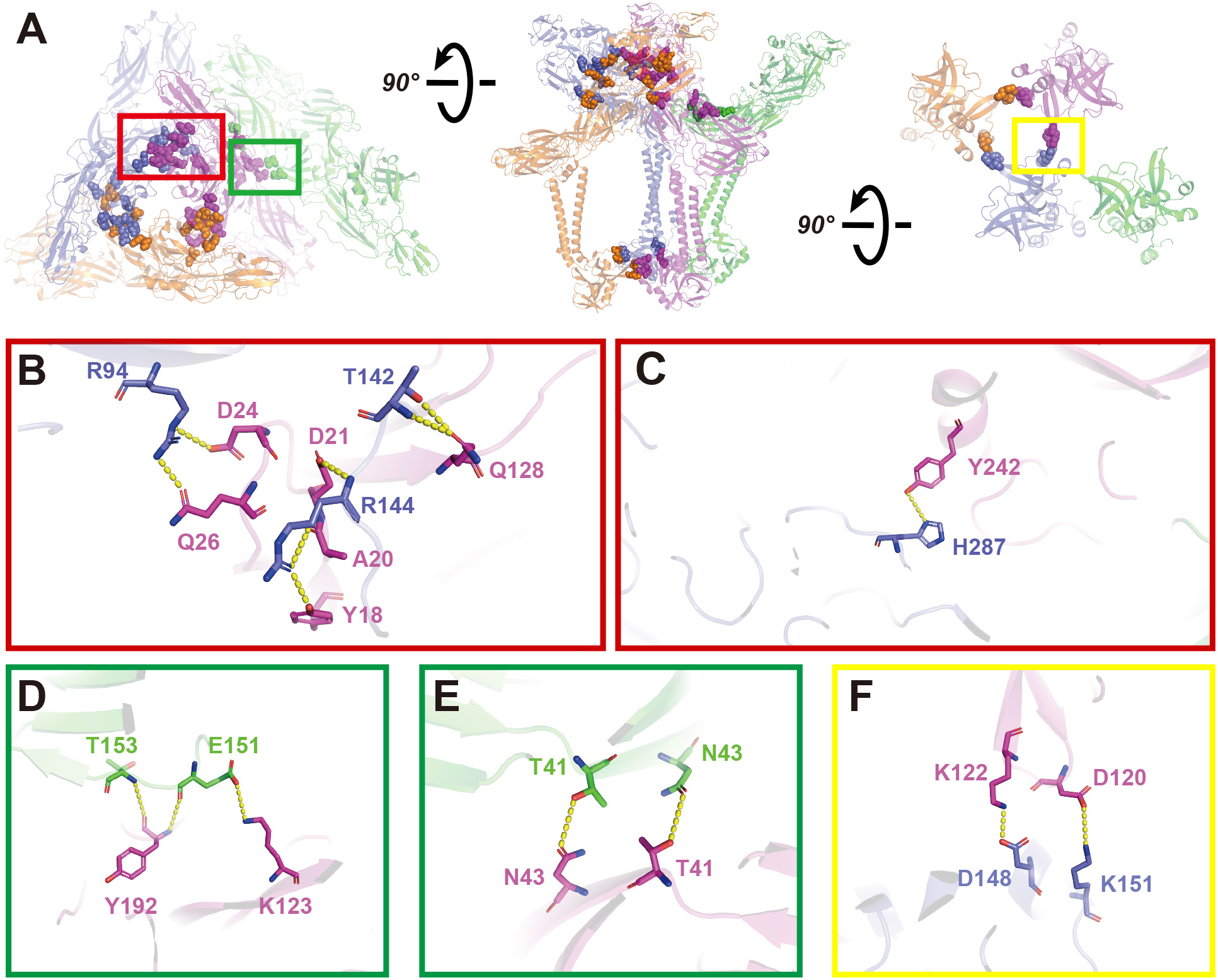
Interaction of GETV protein domains in the ASU. **A**, The top, side, and bottom view of the asymmetric unit (ASU). Four E1-E2-capsid heterotrimers in the ASU are colored slate, magenta, orange, and green. Residues involved in the interaction are displayed as spheres with the same color of the corresponding heterotrimers. The slate, magenta, and orange heterotrimers form the Q-trimer, and the green heterotrimer from neighbor I-trimer. **B**, Zoomed-in view from the red box in A of the interaction between E2 (magenta) and the neighboring E2 (slate) protein. **C**, Zoomed-in view from the orange box in A of the interaction between E1 and the neighboring E2 proteins. **D** and **E**, Zoomed-in views from the green and blue boxes in A, showing the interactions between E1 (magenta) and the neighboring E1 (green), which also represents the interaction between Q-trimer and I-trimer. **F**, The interaction between one capsid protein (magenta) and a neighboring capsid protein (slate) in the Q-trimer. The yellow dashed lines indicate the distance between the atoms involved in the interaction (the cut off distance is 3.5Å).

There are six regions that form contacts between two ASUs (**Figure 4A**). The E1 subunit of Q-trimer from ASU1 interacts with an adjacent Q-trimer from AUS2 in three regions (**Figures 4B, 4C, and 4D**). The first interaction region is located between E1’s domain I in ASU1 and E1’s domain II in ASU2. Multiple residues, including K123, Y129, Y214 and E151, T153, R160 are involved (**Figure 4B**). The second interaction region is located between one E1’s domain I (residues R21, N22, and R289) and another E1’s domain III (residues D385, P382, T307, and D311) (**Figure 4C**). And the third interaction region is located within two E1 proteins, residues D323 and K351 distribute in the corresponding domain III (**Figure 4D**). Residues R437 and A417 are located between the tail of E1 and C-terminal helix of E2 (**Figure 4G**). Except those interaction between the envelope glycoproteins, two sets of capsid-capsid interactions between two ASUs are also observed (**Figures 4E** and **4F**). Interestingly, most of those related interacting residues pairs occur between acidic and basic residues. For instance, E151 and K123 (**Figure 4B**), R21 and D385, R289 and D311 (**Figure 4C**), D323 and K351 (**Figure 4D**), E191 and R230 (**Figure 4E**), K179 and E263 (**Figure 4F**). The electrostatic attractions increase the protein-protein interaction stability.

**Figure 4.**
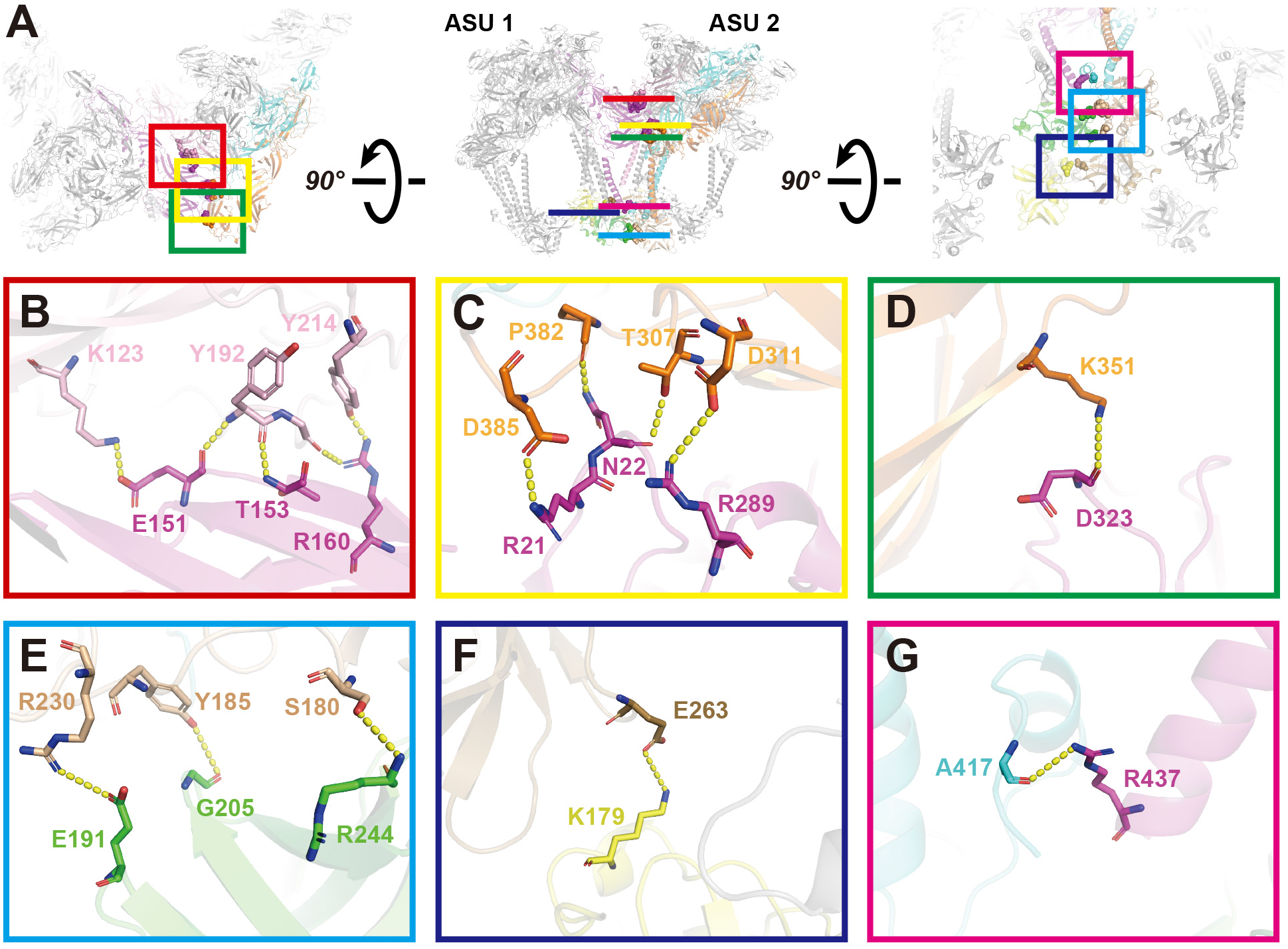
Interaction of GETV protein domains between two ASUs. **A**, The top, side, and bottom view of two adjacent Q-trimers. The E1, E2 and capsid proteins participating in the interaction are colored; other proteins are gray. Residues involved in the interaction are displayed as spheres, in the same color of the corresponding proteins. **B, C**, and **D**, Zoom-in views of the interaction between E1 (magenta) of ASU1 and E1 (pink) of the neighboring E1-E2-capsid heterotrimer, along with another E1 (orange) of the heterotrimer in ASU2. **E** and **F**, Two sets of capsid-capsid interactions occurring between the contiguous capsids of ASU1 and ASU2. Green and yellow capsids are in the ASU1, and the other two capsids are in the ASU2. **G**, Interaction region between E1(magenta, the same E1 protein with **B, C** and **D**) and E2 (cyan, the same heterotrimer with E1 in **C** and **D**). The yellow dashed lines indicate the distance between the atoms involved in the interaction.

We summarize all of the protein-protein interactions in **Table S2**, including hydrogen bonds, salt bridges and van der Waals contacts. There are a total of 47 interactions, among which 28 have been described in earlier studies, and 19 are newly identified in the present study. Taken together, the protein-protein interactions inside E1-E2-capsid heterotrimer, within one ASU, and between ASUs stabilize the structural assembly of the GETV, as well as other alphaviruses.

### Glycosylation sites in the E1-E2 ectodomain are surface-exposed

Viral envelope proteins have evolved to be extensively glycosylated with versatile functions ranging from immune evasion by glycan shielding to enhancement of cell infection (Watanabe et al., 2019). Glycosylation sites have been reported in CHIKV (Voss et al., 2010), EEEV (Hasan et al., 2018), SINV (Chen et al., 2018), and MAYV (Ribeiro-Filho et al., 2021). We first predicted the N-glycosylation and O-glycosylation sites on GETV E1/E2 using NetNGlyc (Gupta and Brunak, 2002), NetOGlyc (Steentoft et al., 2013), GlycoMine (Li et al., 2015), and YinOYang (Gupta and Brunak, 2002). A list of the predicted glycosylation sites is provided in **Table S3**. We next utilized a mass spectrometry method to identify glycosylation sites in the GETV E1 and E2 glycoproteins a list of the detected sites is given in **Table S3**. In the 2.8 Å-resolution cryo-EM density map, we sought the densities next to the residues which were not assigned to proteins. A total of eight densities were observed: four in E1, including S66, N141, N270, and N335. Another four in E2, T155, N200, T264, and N262 (**Figure 5**). Four densities for glycans were modeled and fitted associated with N141 and N270 of E1, and N200 and N262 of E1 (**Figures 5D-5G**). Although protruding densities adjacent to S66, T155, T264, and N335 are strong, these did not support precise fitting for glycan atomic models (**Figures 5H-5K**).

**Figure 5.**
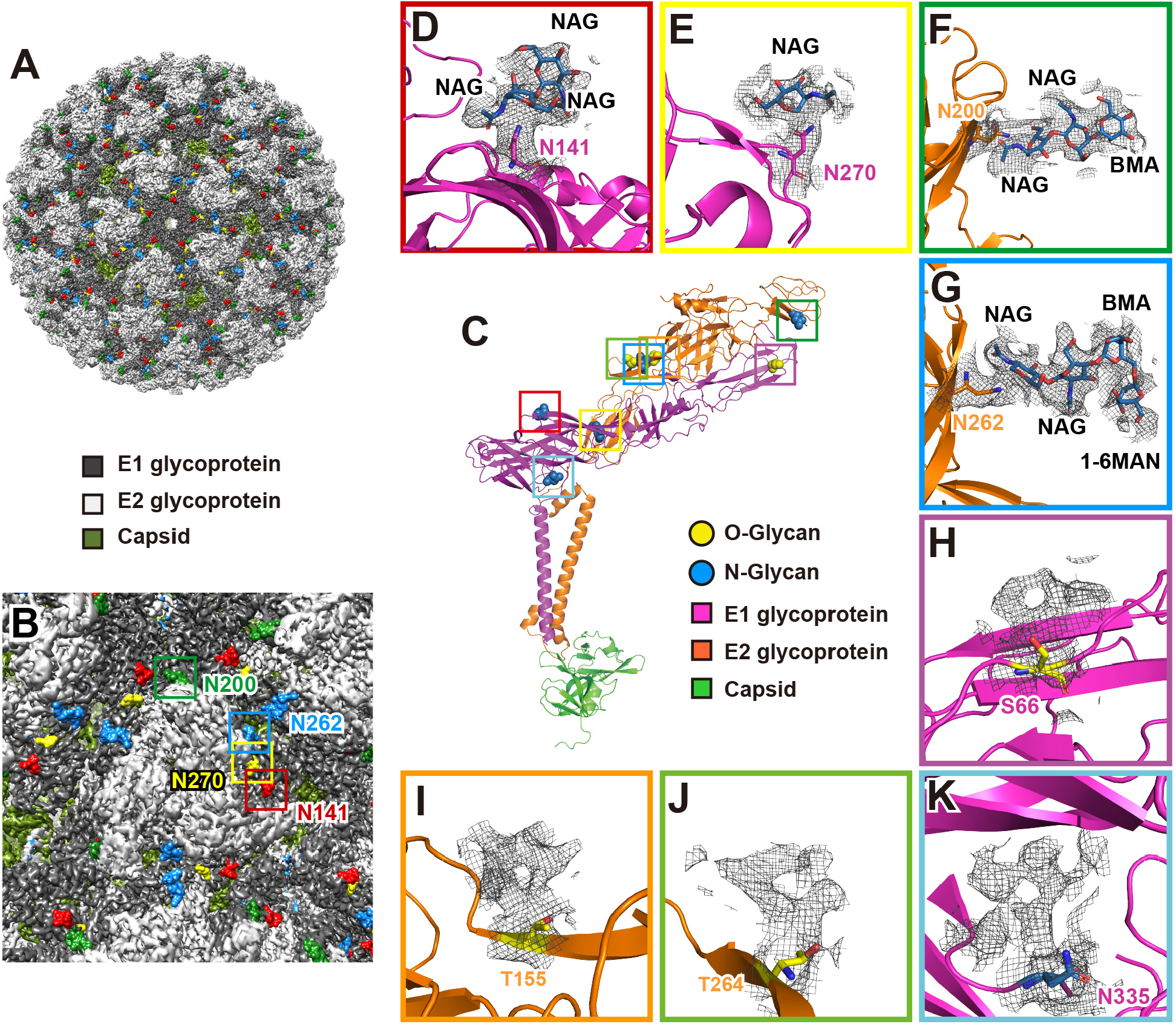
Atomic models of glycans fitted in the density maps in the E1-E2 ectodomain. **A**, External surface of a GETV virion, with glycosylation sites displayed in color (E1 N141 in red, E1 N270 in yellow, E2 N200 in green, and E2 N262 in blue, respectively). **B**, Zoomed-in of the external surface close to the icosahedral 3-fold vertices; the glycosylation sites are framed and labeled using the same color scheme as shown in panel A. **C**, Glycosylation sites in the atomic model of E1-E2-Capsid heterotrimer. **D** – **G**, Zoomed-in views of N-glycosylation sites with residues shown as sticks (N141 and N270 in E1 are in purple, and N200 and N262 in E2 in orange). Cryo-EM densities attributed to glycans are shown as a mesh; the atomic models for the glycans are shown as blue sticks. The names of carbohydrate monomers are labeled. N-Acetyl-glucosamine (NAG) and mannose (BMA: β-D-mannose; 1-6MAN: α- (1-6)-D-mannose) were assigned as the hexosamine and hexose monomers based on cryo-EM density map interpretation and models previously built for MAYV (Ribeiro-Filho et al., 2021). **H** – **K**, Zoomed-in views of glycosylation sites with residues shown as sticks, cryo-EM densities attributed to glycans are shown in mesh. Although protruding densities adjacent to S66, T155, T264, and N335 are strong, these did not support precise fitting for glycan atomic models.

The N-glycosylation of E1 N141 and E2 N262 have also been reported in the MAYV structure, for which glycan models were built (Ribeiro-Filho et al., 2021). In our GETV structure, densities associated with E1 N141 and E2 N262 were both clearly visible, allowing fitting of atomic models for multiple hexosamine and hexose monomers. The proximal part of the density associated with E2 N262 were accommodated three carbohydrates, which were modeled as classical N-glycans NAG-NAG-BMA (NAG: N-Acetyl-glucosamine and BMA: β-D-mannose, **Figure 5G**). At the position of the third carbohydrate BMA, one branch density was observed, in which a 1-6MAN (α- (1-6)-D-mannose) monomer was modeled and fitted (**Figure 5G**). A similar model of NAG-NAG was fitted in the density proximal to E1 N141 (**Figure 5D**). E2 N200 has a density that also fitted the typical N-glycans, NAG-NAG-BMA (**Figure 5F**). The E1 N270 has a less defined density compared to E1 N141, E2 N200 or N262: only one NAG monomer could be modeled here (**Figure 5E**). All four glycan moieties are surface-exposed (**Figures 5A** and **5B**), they are accessible on the GETV surface close to the trimer/trimer interface (**Figure 5B**). This surface exposure of glycosylation sites suggests their potential roles in evading detection by the host immune system, and/or in promoting attachment to the host cells to enhance viraluptake.

### S-acylation sites increase the stability of E1 and E2 TM helixes

S-acylation is a post-translational modification that attaches fatty acids, often palmitic acid, to cysteine residues; this modification occurs on both peripheral and integral membrane proteins (Linder and Deschenes, 2007). These fatty acids increases a protein’s hydrophobicity and also facilitates interactions with lipid bilayers, and S-acylation is known to functions directly in modulating proteins’ stability (Linder and Deschenes, 2007). Protein acylation was originally discovered with vesicular stomatitis virus (Schmidt and Schlesinger, 1979). Over the past 40 years, numerous viruses have been reported to undergoing S-acylation, such as human immunodeficiency virus (HIV), severe acute respiratory syndrome coronavirus (SARS-CoV), influenza A virus, and hepatitis C virus (HCV) (Veit, 2012). For alphaviruses, both SFV and SINV have been reported to contain S-acylation sites (Ivanova and Schlesinger, 1993; Ryan et al., 1998; Veit et al., 1996; Zhao et al., 1994). In the cryo-EM structure of BFV at 6 Å-resolution, density has been found next to residue C394 in the E2 TM helix; however, no atomic model of a fatty acid was built (Kostyuchenko et al., 2011).

In the present study, we first predicted the possible S-acylation sites using predictors CSS-Palm and GPS-Palm (Ning et al., 2021; Ren et al., 2008). A list of predicted S-acylation sites is given in **Table S4**. Next, we utilized a mass spectrometry method to identify S-acylation sites in the GETV E1 and E2 proteins, and list of the sites are stated in **Table S4**. Finally, in the 2.8 Å-resolution GETV map, we examined the densities next to the residues. A total of five such densities have been identified: one at E1 C433, and four in E2 (C385, C395, C415 and C416) (**Figure 6**). According to the mass spectrometry analysis, stearic acids are attached to C385 and C395 in E2, while other three cystines, C433 in E1 and C415, C416 in E2, are acylated with palmitic acids. Therefore, we accommodated atomic models of stearic acid in the straight proximal portion of the density next to E2 C385 and C395 (**Figure 6C** and **6D**), and fitted atomic models of palmitic acid in the densities adjacent to E1 C433 and E2 C415 and C416 (**Figures 6B, 6E**, and **6F**). As illustrated in **Figure 6A**, the five high or absolutely conserved C-terminal S-acylation sites strongly anchored the E1 and E2 TM helixes into the inner leaflet of lipid bilayer of viralenvelop membrane. Considering the large portion of both E1 and E2 ectodomain (**Figure 1E**), S-acylation in the TM helixes appears likely to stabilize the E1/E2 glycoproteins.

**Figure 6.**
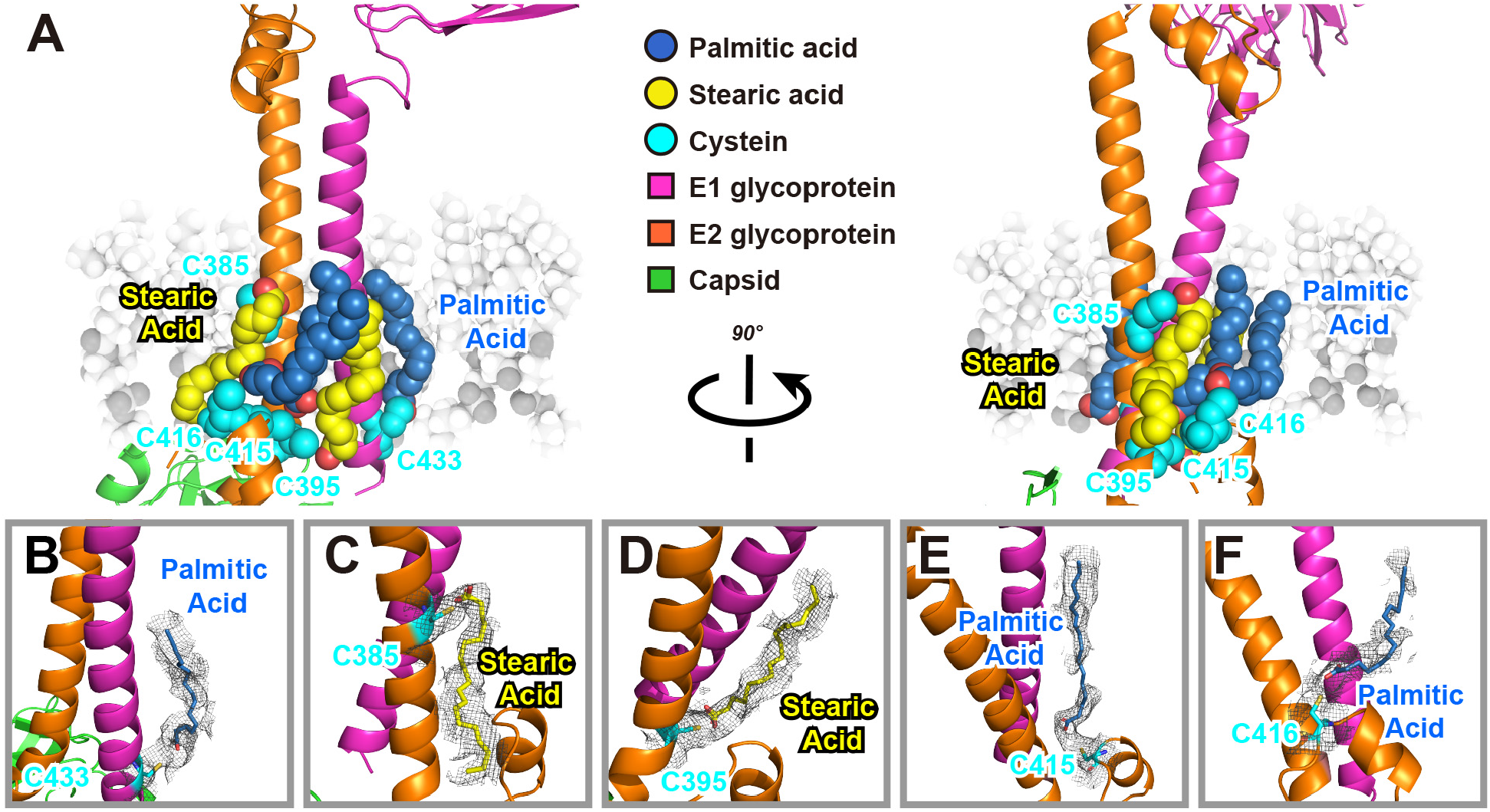
S-acylation sites identified in the E1 and E2 TM helixes and in the E2 cytoplasmic tail. **A**, Overview of S-acylation sites of E1 and E2 proteins. E1 and E2 are presented as cartoons (in Pymol), and are colored using the same color scheme as in Figure 2. The involved cysteines, palmitic acids, and stearic acids are presented as spheres, colored with cyan, steal blue, and yellow. The inner leaflet of viralenvelop membrane is shown in the background. **B** – **F**, Zoom-in views of S-acylation sites. Palmitic acids and stearic acids are shown as sticks. Cryo-EM densities attributed to the cysteines, as well as palmitic acids and stearic acid, are shown as a mesh.

### Cholesterol molecules and a phospholipid molecule in the hydrophobic pocket increase stability of E1-E2 heterodimer

Previous studies have demonstrated a hydrophobic pocket is essential for alphavirus assemble at the outer leaflet of the envelop membrane (Chen et al., 2018; Ribeiro-Filho et al., 2021). The hydrophobic pocket comprises the TM helix from E1, and the TM helix and domain D of E2 (see **Figure 1E**). In the 3.5 Å-resolution SINV structure, a “pocket factor” was identified; this was predicted to be a hydrophobic phospholipid tail (Chen et al., 2018). In the 4.4 Å-resolution MAYV structure, a long extra density was identified in the hydrophobic pocket, and a C18 hydrocarbon lipid molecule, octadecane, was fitted into the density (Ribeiro-Filho et al., 2021).

In our GETV structure, extra densities were clearly visible in the cavity formed by the E1 and E2 TM helixes, as well as in the region immediately adjacent to the helixes (**Figures 7A– 7C**). Two cholesterols were modeled and fitted into the densities adjacent to the helixes (**Figure 7B**), while one cholesterol was fitted in one density in the cavity (**Figure 7C**). Cholesterol is the most abundant sterol in mammals, it is a major constituent of the plasma membrane and plasma lipoproteins. A previous study revealed that nearly 30% of the lipid class composition of baby hamster kidney cells (BHK, one of the most commonly used cell lines for expression of alphaviruses, and GETV in the present study) represents cholesterol, and more than 40% of the envelop membrane of SFV particles produced in BHK cells are cholesterol (Kalvodova et al., 2009). Another density in the cavity displayed a “Y” shape, and a phospholipid molecule was modeled as “DOPC” (dioleoyl-phosphatidylcholine, **Figure 7C**), with a hydrophilic “head” containing a phosphatidylcholine, and two hydrophobic “tails” derived from oleic acid, a mono-unsaturated omega-9 fatty acid. Both densities that fitted with two oleic acid hydrocarbon tails have a kink that can accommodate the mono-unsaturated bond (**Figure 7C**).

**Figure 7.**
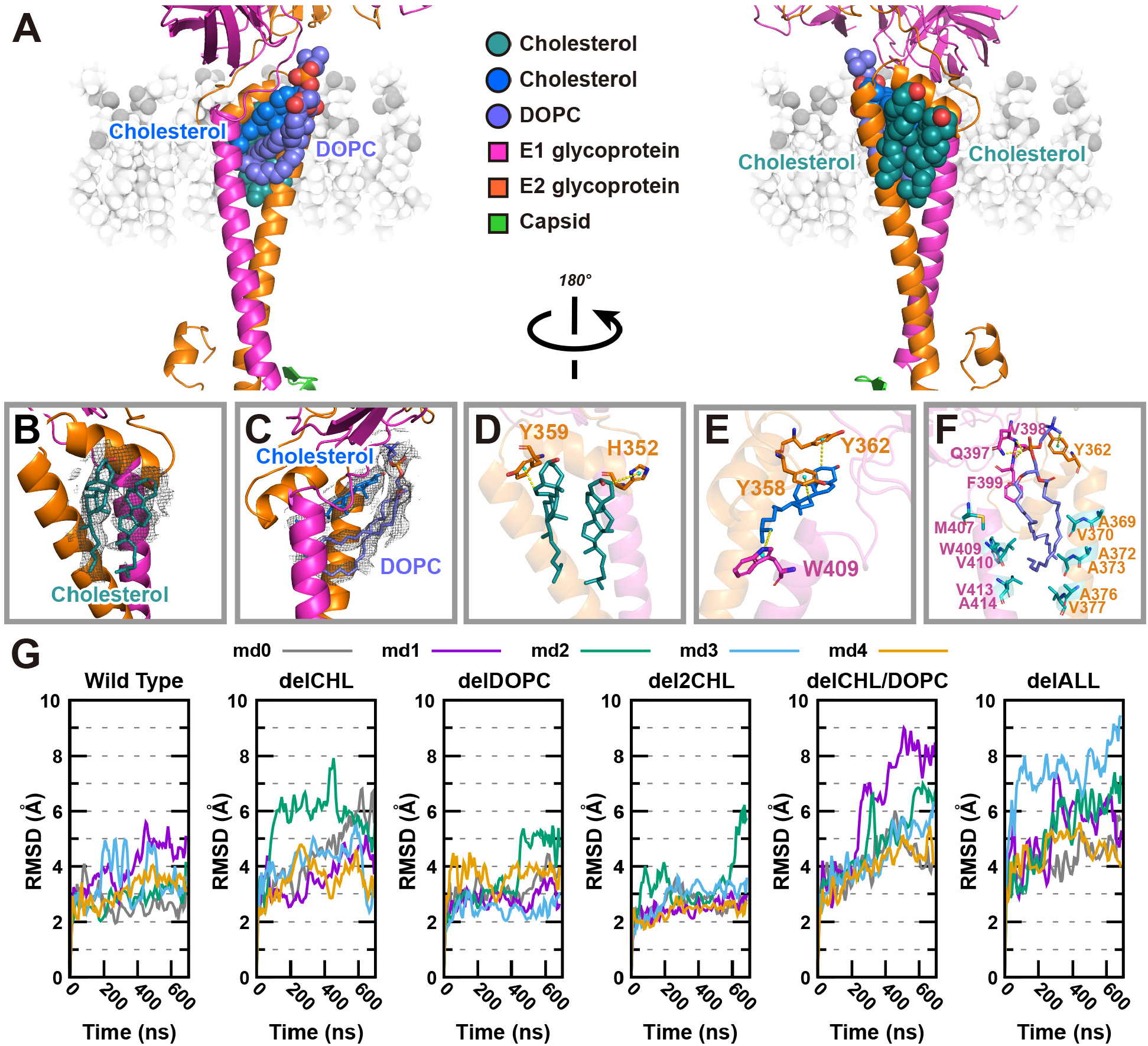
Cholesterol molecules and a phospholipid DOPC in the hydrophobic pocket. **A**, Overview of the distribution of cholesterols and a phospholipid molecule DOPC. A cholesterol (blue) and DOPC (slate) are in the hydrophobic pocket formed by the E1 TM helix, the E2 TM helix, and E2 domain D. The other two cholesterols (deep teal) are positioned on the other side of domain D. E1 and E2 are presented as cartoons. The involved cholesterols and DOPC are presented as spheres, and outer leaflet of viralenvelop membrane is shown in the background. **B** and **C**, Zoom-in views of cholesterols and DOPC molecules, which are shown as sticks. Cryo-EM densities attributed to the cholesterols and phospholipid are shown as a mesh. **D** – **F**, Zoom-in views of the interactions between cholesterol or DOPC and E1 or E2. The aromatic residues positioned near the cholesterol or DOPC are shown as sticks with the same color as E1 or E2. The hydrophobic residues positioned near the tail of DOPC are shown as sticks. The center of mass of aromatic ring are shown as nb_spheres, colored in cyan. The yellow dashed lines represent the hydrogen bonds. A portion of the hydrogen bonds are CH-π hydrogen bonds between the soft acids CH and a soft base π-electron system. The remaining portion are common hydrogen bonds between the phosphate group of DOPC and Q397, V398 or F399. **G**, Time evolutions from MD simulation analysis for the Cα-RMSD of E1 (295-435) and E2 (269-422) for systems “Wild Type”, “delCHL”, “delDOPC”, “del2CHL”, “delCHL/DOPC”, and “delALL”. All curves were smoothed with the Bezier method, implemented in gnuplot.

We subsequently analyzed the interactions between E1/E2 and cholesterol/DOPC in the hydrophobic pocket. Cholesterol molecules contain four fused carbon rings referred to as A, B, C and D. It is the principal lipid interact with proteins through CH-π hydrogen bonds. CH-π bonds involving a soft acid CH and a soft base π-electron system (Nishio et al., 2014). As shown in **Figure 7D**, in the cholesterol at left, the B ring interacted with aromatic ring from Tyr-359 in subdomain D, with mean distances of 4.0 Å. The A ring in the right cholesterol contacted with aromatic ring from His-352, with a distance of 4.4 Å. This is consistent with the establishment of the cut-off values (4.5 Å) for the distance of the donor carbon atoms to the geometric center of the π-acceptor system (Nishio et al., 2014). **Figure 7E** displayed the cholesterol in the hydrophobic cavity, the A and D rings interacted with Tyr-362 and Tyr-358, with mean distances of 4.3 Å and 3.8 Å, respectively. A CH-π hydrogen bond between a methyl group of DOPC and aromatic ring of Tyr-362 were identified, with a distance of 4.5 Å. Moreover, hydrogen bonds between the phosphate group and Gln-397/Val-398/Phe-399 were confirmed. Beyond that, multiple hydrophobic residues were found around hydrophobic tails of DOPC. All of these interactions fixed DOPC in the hydrophobic pocket (**Figure 7F**).

To further investigate how the cholesterol and DOPC molecules regulate the E1-E2 complex, we calculated the root mean square deviation of the Cα atom (Cα-RMSD) of E1-E2 against time in the six simulated systems using molecular dynamics (MD) simulations (**Movie S2**). As shown in **Figure 7G**, the RMSD evolves differently in the six systems, with the overall values from the “delCHL/DOPC” (both cholesterol and DOPC in the pocket were removed) and the “delALL” (all 3 cholesterols and DOPC were removed) the most unstable. The RMSD values increase drastically higher in these two systems with the magnitude reaching as high as 9 Å. Whereas those in the “delCHL” (only cholesterol in the pocket was removed) system, the “delDOPC (only DOPC in the pocket was removed) system, and the “del2CHL” (2 cholesterols surround the pocket were removed) system populate similarly to or only slightly higher than in the Wild Type system. This indicates that the structure of E1-E2 complex changes significantly whenever the pocket cholesterol and DOPC ligands are removed together, suggesting a cooperative role between the two pocket ligands in stabilizing the E1-E2 complex.

## Discussion

Viral glycosylation has broad roles in viral pathobiology, including mediating protein folding and trafficking, facilitating viral budding and release, directing immune evasion by glycan shielding, and shaping viral infection and tropism (Watanabe et al., 2019). For alphaviruses, glycosylation is known to be a key determinant of cellular tropism and virulence: a lack of glycosylation in RRV rendered virions unable to induce alpha/beta interferon production in in myeloid dendritic cells, the professional antigen-presenting cells that rapidly respond to viruses, suggesting that viral glycans may negatively regulate antiviral responses (Shabman et al., 2008). Glycosylation has also been demonstrated to function in alphavirus infection and replication: a lack of glycosylation for the E1 protein in SINV severely impaired viral replication (Knight et al., 2009). Further, a study showed that mutations of the glycosylation sites in the Salmonid alphavirus E1 and E2 proteins severely altered the virulence and production of an infectious virus (Aksnes et al., 2020).

Previous structural analyses have revealed glycosylation sites in alphaviruses. A crystal structure of the mature E3-E2-E1 glycoprotein complexes from CHIKV revealed three N-linked glycans: N12 in E3, N263 in E2, and N141 in E1 (Voss et al., 2010). The cryo-EM structure of MAYV identified two glycosylation sites: N141 in E1 and N262 in E2, with at least five carbohydrates modeled as NAG-NAG-BMA-MAN-MAN at the E2 N262 site and with the canonical NAG-NAG-BMA glycan sequence modeled at the E1 N141 site (Ribeiro-Filho et al., 2021). Both the E1 N141 and E2 N262 sites are conserved in CHIKV, MAYV, SFV, RRV and GETV. Glycosylation sites were also identified in SINV by cryo-EM, including N139 and N245 in E1, N283 in E2, and N14 in E3, all of which displayed extra density next to asparagine residue, although no glycan models were built (Chen et al., 2018). In the present study, six new glycosylated residues – E1 S66, N270, N335 and E2 T155, T264, and N200 – were identified, and we built two more atomic models of the glycans (N270 and N200) precisely into the 2.8 Å cryo-EM density map (**Figure 5**).

Protein S-acylation is a post-translational lipid modification wherein fatty acids, usually palmitic acid, are attached to cysteine (S-palmitoylation). There are also know examples of modifications of serine and threonine residues (O-palmitoylation) (Veit, 2012). S-acylated proteins are typically membrane proteins, S-acylation enhances protein hydrophobicity and impacts protein subcellular localization, trafficking, stability, and interactions with other proteins (Wu et al., 2021). S-acylation of viral proteins is known to be involved in virus assembly and infection. For example, the Spike protein of severe acute respiratory syndrome coronavirus (SARS-CoV), and quite recently, of SARS-CoV-2, both have been shown to be palmitoylated; and these modifications have been associated with the cell-cell fusion process known to be essential for viral infectivity (McBride and Machamer, 2010; Wu et al., 2021). For alphaviruses, both E1 and E2 glycoproteins of SFV have been reported to contain S-acylation sites: E1 contains mainly stearic acid, while E2 is acylated primarily with palmitic acid (Veit et al., 1996). Palmitoylation of cysteine residue in the E2 protein of SFV was demonstrated as an essential determinant for viral budding (Zhao et al., 1994). The E1 and E2 protein of SINV also have palmitoylation sites, and site-directed mutagenesis of cysteines in E1 and E2 abolished fatty acids attachment, and result in aberrant viral assembly and particle formation, slow replication at early times post-infection, and increased sensitivity to detergents (Ivanova and Schlesinger, 1993; Ryan et al., 1998). Those findings supported an earlier study which indicated that mutations in the SINV E2 glycoprotein led to defects in palmitic acid attachment, as well as defects in virus assembly and budding (Gaedigk-Nitschko and Schlesinger, 1991). In the cryo-EM structure determination of BFV at 6 Å-resolution, density was detected next to one cystine residue in the E2 protein (C394); however, no atomic model of a fatty acid was built (Kostyuchenko et al., 2011). In another cryo-EM study of VEEV at 4.4 Å-resolution, three conserved E2 cysteine residues (C396, C416, and C417) were mapped near the lipid head groups of the inner membrane; note that no density attributable to S-acylated fatty acids was found (Zhang et al., 2011).

Cholesterol is an essential structural component of cell membranes and can also serve as a structural component of viral envelop membranes. The membrane of mammalian-derived SFV virions comprises more than 40% cholesterol (Kalvodova et al., 2009). Interestingly, the cholesterol content in the envelope of MAYV particles obtained from mammalian cells is 10-fold higher in cholesterol content as compared to viruses produced in mosquito cells (Sousa et al., 2011). A similar discrepancy in cholesterol content was found in SINV grown in mammalian vs in insect cells (Hafer et al., 2009). In all studies, BHK cells were used as the mammalian host cells; these are the same cells we used in the present study to propagate and purify GETV. Thus, it was not surprising that cholesterol was present in the GETV envelop membrane. It was however surprising that we detected one cholesterol molecule in the hydrophobic pocket and two cholesterol molecules surrounding the TM helixes of the E1 and E2 proteins, which indicated that cholesterol interacts with the E1 and E2 proteins (**Figure 7**).

Several studies have addressed the potential impacts of cholesterol present in viral envelope on viral infections, and depletion of cholesterol from envelopes has been shown to abolish the infectivity of influenza virus, human herpesvirus, herpes simplex virus, hepatitis C virus, and HIV (Aizaki et al., 2008; Campbell et al., 2002; Huang et al., 2006; Sun and Whittaker, 2003). For alphaviruses CHIKV, SFV, SINV, and MAYV, cholesterol is required for viral entry to host cells (Carvalho et al., 2017; Lu et al., 1999; Phalen and Kielian, 1991; Tsetsarkin et al., 2007). After entry, viral RNA is released and replicated; this process also depends on intracellular membranes, and cholesterol impacts on viral replication: SINV replication was faster and reached higher titer in cholesterol-rich fibroblasts, and virions produced in cholesterol-rich fibroblasts are more infectious than viral particles produced from normal human fibroblasts (Ng et al., 2008). Similarly, drugs that induced intracellular cholesterol accumulation and affected cholesterol biosynthesis, conferred strong inhibition against CHIKV replication (Wichit et al., 2017). Notably, gene expression profiling of cells infected with M1 strain of GETV (which was isolated from *Culex* mosquitoes and known to infect horses and pigs (Zhai et al., 2008)), demonstrated that more than 60% of genes associated with the cholesterol biosynthesis pathway are downregulated after GETV M1 infection (Liang et al., 2018). Taken together, cholesterol is not only a molecular factor that essential for alphavirus entry, replication, budding, and exit from host cells, but also a structural factor contributing to the stability of viral hydrophobic pocket in the envelop membrane.

## STAR METHODS

Detailed methods are provided in the online version of this paper and include the following:

- KEY RESOURCES TABLE
- RESOURCE AVAILABILITY
  - Lead Contact
  - Materials Availability
  - Data Availability
- METHOD DETAILS
  - Virus Infection
  - Virus production and purification
  - Cryo-EM sample preparation and data acquisition
  - Image processing
  - Model building and refinement
  - Quantitative N-glycosylation analysis by LC-MS/MS
  - Quantitative S-acylation analysis by LC-MS/MS
  - Molecular dynamics simulation study of cholesterols and DOPC regulation of E1-E2

## Supporting information

Star Methods

## Supplemental Information

Supplemental Information included 7 figures, 4 tables, and 2 movies.

## Acknowledgements

We thank all staff members of the Cryo-electron Microscopy Centre, Southern University of Science and Technology, for their assistance with data collection. This work was supported by National Natural Science Foundation of China (81870246 and 82070329 to Z.L.), and by Guangdong Basic and Applied Basic Research Foundation (2019A1515110090 to R.Q.).

## Author contributions

Conceptualization, Z.L. and C.W.; Methodology, Z.L., C.W., C.L., and A.W.; Investigation, A.W, F.Z., C.L., D.G., R.Q., Y.Y., S.L., Y.G., L.F., and Y.X.; Resources, Z.L., Y.X., and C.W.; Writing – Original Draft, Z.L. A.W., C.L., and C.W.; Writing – Review & Editing, Z.L.; Supervision, Z.L. and C.W.; Funding Acquisition, Z.L.

