## Supplemental Figures and Tables for "Structure of Infective Getah Virus at 2.8 Å-resolution Determined by Cryo-EM"

### Supplemental Information

**Figure S1.** GETV is a mosquito-borne arbovirus that caused reproductive disorders in pregnant mice.

**Figure S2.** Cryo-EM image-processing workflow.

**Figure S3.** Local resolution of cryo-EM block-based reconstructions of GETV and density maps of structure domains from E1 and E2 proteins.

**Figure S4.** Density maps of capsid protein.

**Figure S5.** Multiple sequence alignment and secondary structural elements of alphavirus E1 protein.

**Figure S6.** Multiple sequence alignment and secondary structural elements of alphavirus E2 protein.

**Figure S7.** Multiple sequence alignment and secondary structural elements of alphavirus capsid protein.

**Table S1.** Cryo-EM data collection and processing, block-based reconstruction, model building and refinement statistics.
**Table S2.** Protein-protein interactions in GETV and other alphaviruses.
**Table S3.** Glycosylation sites in E1 and E2 proteins.
**Table S4.** S-acylation sites in E1 and E2 proteins.

**Movie S1.** GETV caused mobility impairments in pelvic limbs in newborn mice.
**Movie S2.** Molecular dynamics simulation of cholesterols and DOPC interaction with E1-E2.

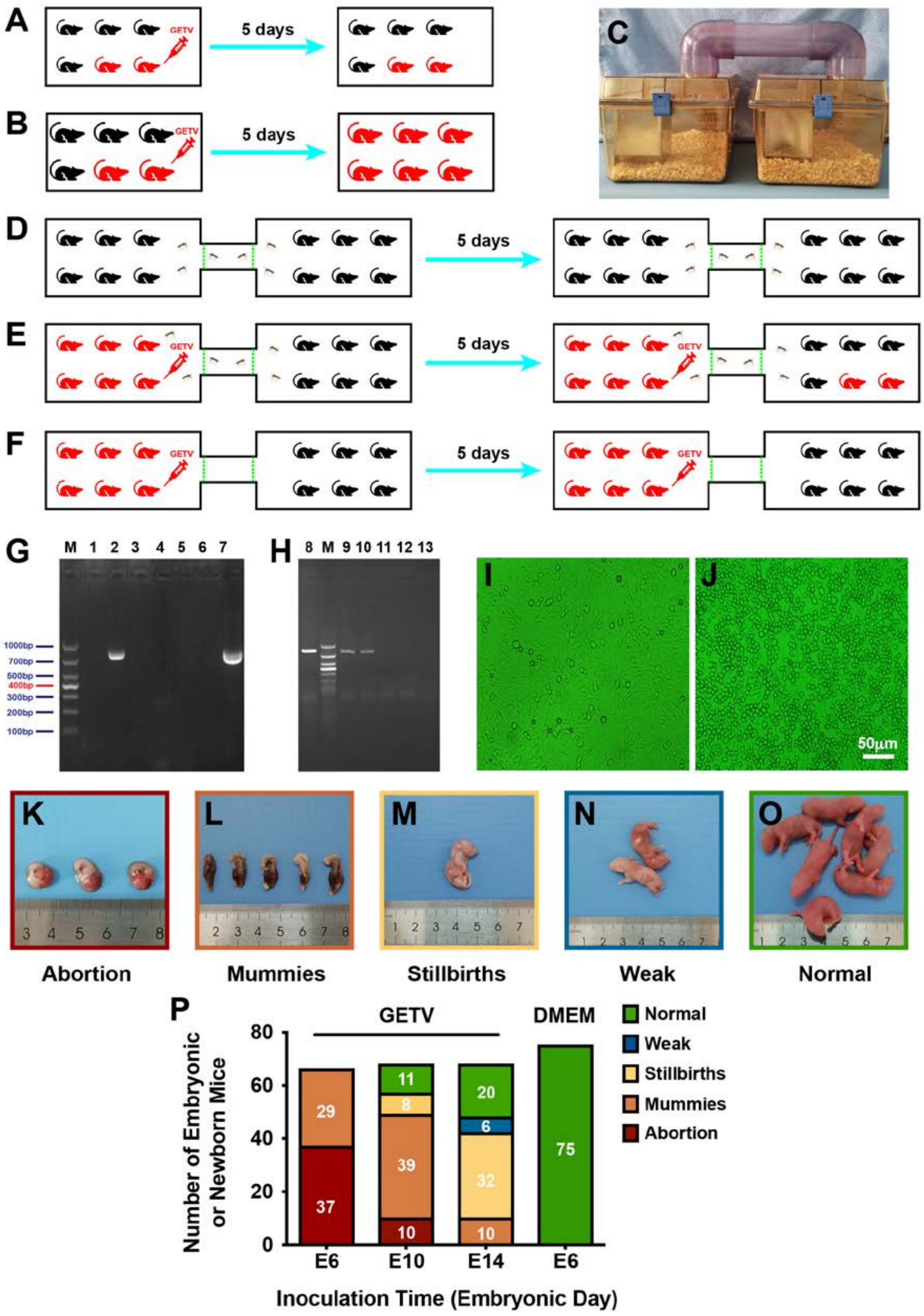


**Figure S1. GETV is a mosquito-borne arbovirus that caused reproductive disorders in pregnant mice.**

**A** and **B**, **D**–**F**, schematic illustrations of the experimental design to investigate the infectiousness of GETV. **C**, photo of the device that was used in experiments **D**–**F**. **G** and **H**, Total RNA was extracted from tissue, including spleen, lung, cerebral cortex, and various lymph nodes using TRIzol reagent after 5 days post inoculation (DPI) (Kumanomido et al., 1988), and subjected to RT-PCR for GETV detection using specific primers (GETV-F: 5′-ACCGAAGAAGCCGAAGAA-3′, and GETV-R: 5′-GCACTCRAGGTCATACTTG-3′) (Zhou et al., 2020). M, markers; Lane 1, buffer used in RT-PCR; Lane 2, supernatant of GETV cultured in BHK21 cells (positive control); Lane 3, test in mosquitos used in D; Lane 4 and 5, test in mice in the right-hand cage at 5 DPI in D; Lanes 6 and 7, GETV-free mice and infected mice in the cage at 5 DPI in A; Lane 8, GETV-free mice at 5 DPI in B, showing GETV positive, indicating the GETV could be transmitted via animal bite. Lanes 9 and 10, GETV-free mice in the right-hand cage in E, showing positive and negative at 5 DPI, demonstrating GETV transmitted from mice in the left-hand cage via mosquitos. Lanes 11-13, three mice in the right-hand cage in F tested negative, revealed that GETV is not an airborne virus. **I** and **J**, tissue samples from the GETV-free and GETV-infected mice in right-hand cage in experiment E, cultured in BHK21 cells, images were captured after 35 hours. Morphological characteristics from viral infections obviously appeared, such as enlarged, rounded, refractile, pyknosis and detachment.

Twenty pregnant mice were randomly divided into four groups. The first three groups were inoculated oronasally with 100 ml ($10^6$ $TCID_{50}$/mL) GETV V1 strain at E6 (embryonic day 6, early-gestation), E10 (embryonic day 10, middle-gestation), E14 (embryonic day 14, late-gestation), respectively. The control group was inoculated with an equal volume of DMEM at E6. **K**, abortions (embryonic were expelled from the uterus before parturition), **L**, mummies (embryos were lost before farrowing), and **M**, stillbirths (embryos were lost around the time of birth, and may be prepartum or intrapartum) arising from GETV-infection were dominant in the E6, E10, and E14 groups. Some weak (**N**) and normal (**O**) newborns were found in E10 and E14 groups. **P**, summary of the proportion of embryonic or newborn mice in the four groups. Numbers in the color frames in each column represented the number of mice with each reproductive disorder.

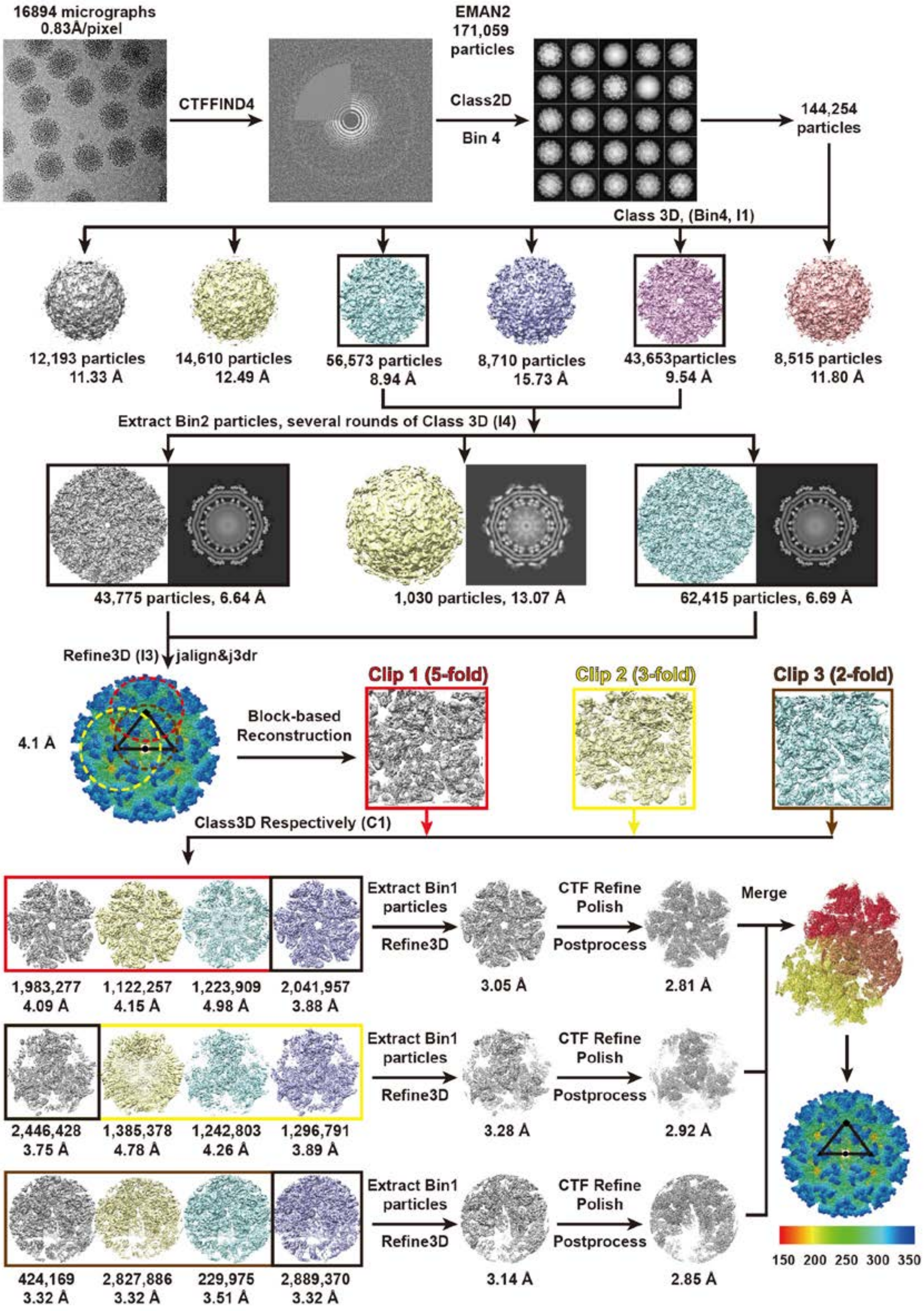


**Figure S2. Cryo-EM image-processing workflow.**

Schematic of pre-processing, 2D and 3D classification, block-based reconstruction, and refinement procedures used to generate the 2.8-Å resolution density map.

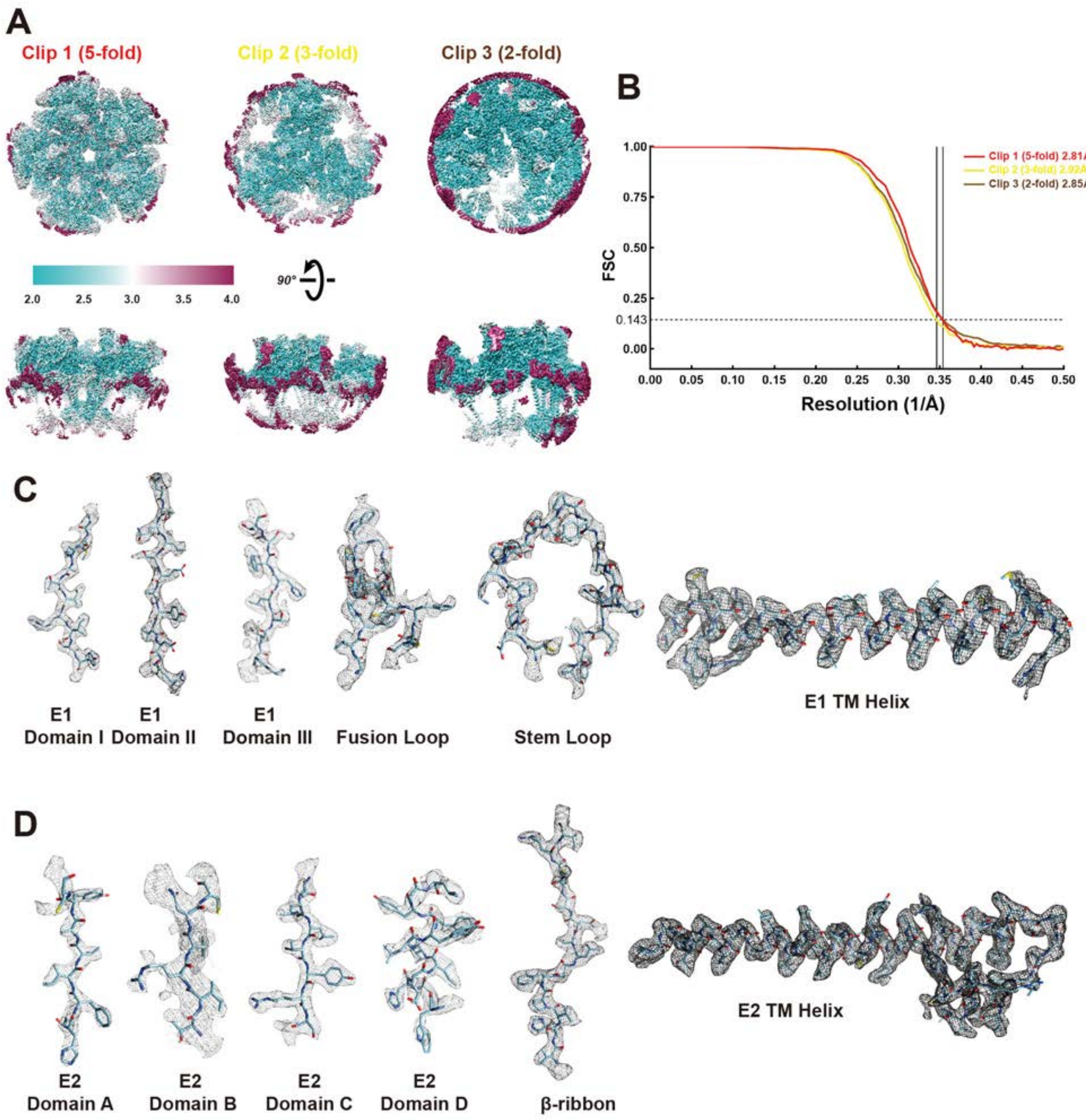


**Figure S3. Local resolution of cryo-EM block-based reconstructions of GETV and density maps of structure domains from E1 and E2 proteins.**

**A**, Block-based reconstructions of GETV, colored according to local resolution. The resolution varies from 2.0 Å (sky blue) through 3.0 Å (light grey) to 4.0 Å (violet red). **B**, Fourier shell correlation (FSC) curve of the final reconstructions indicating an average resolution of 2.81 Å, 2.92 Å, and 2.84 Å for the Clip 1 (5-fold symmetry), Clip 2 (3-fold), and Clip 3 (2-fold), respectively, according to the gold-standard criterion (FSC=0.143). **C**

and **D**, Representative density and atomic models of each domain of E1 and E2 proteins. E1 Domain **I**, residues A128-Y137; E1 Domain **II**, residues N100-R110; E1 Domain **III**, residues K351-T358; Fusion loop, residues V84-T98, Stem loop, residues C380-D401; E1 TM Helix, residues T405-R438; E2 Domain A, residues A92-H99; E2 Domain B, residues T196-C201; E2 Domain C, residues T292-S298; E2 Domain D, residues W350-L361; E2 β-ribbon, K253-V267; E2 TM Helix, residues Y362-R419.

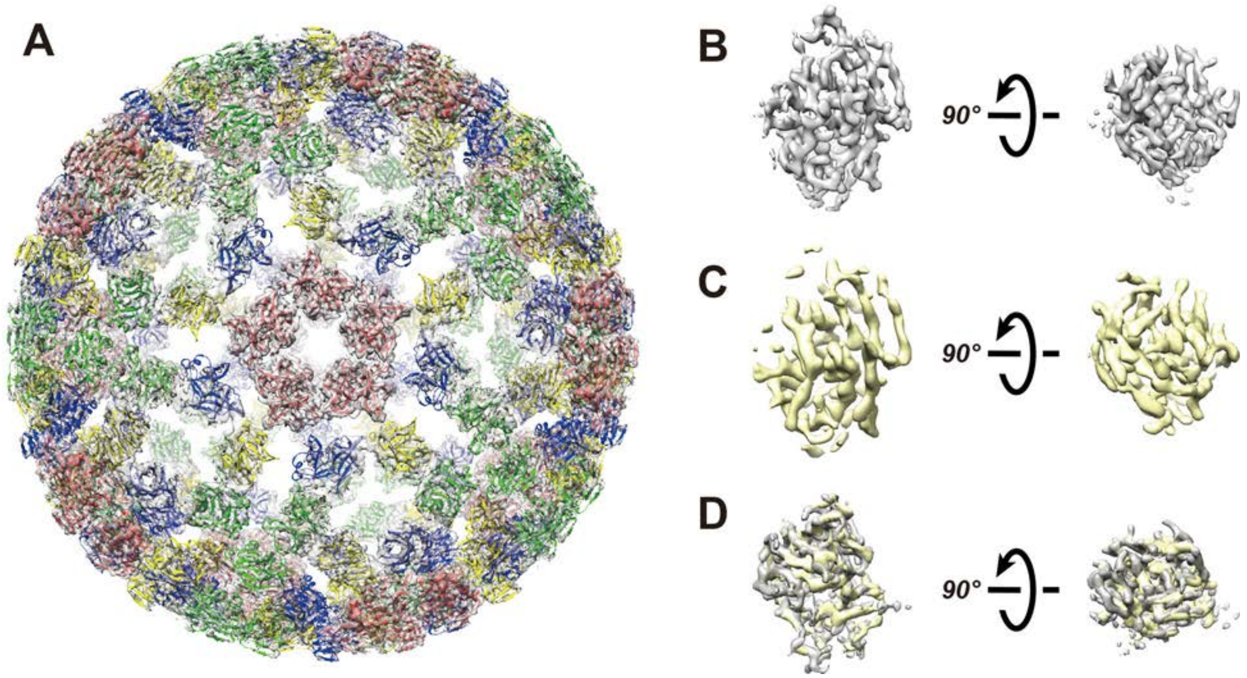


**Figure S4. Density maps of capsid protein.**

**A**, Density map of icosahedral capsid and atomic models. The pentamer capsids models are colored by red, and the other three hexamer capsids models are colored by green, yellow and blue separately. **B**, Density map of the pentamer capsid, **C**, hexamer capsid, and **D**, superpose map of pentamer capsid and hexamer capsid. The correlation value is 0.92.

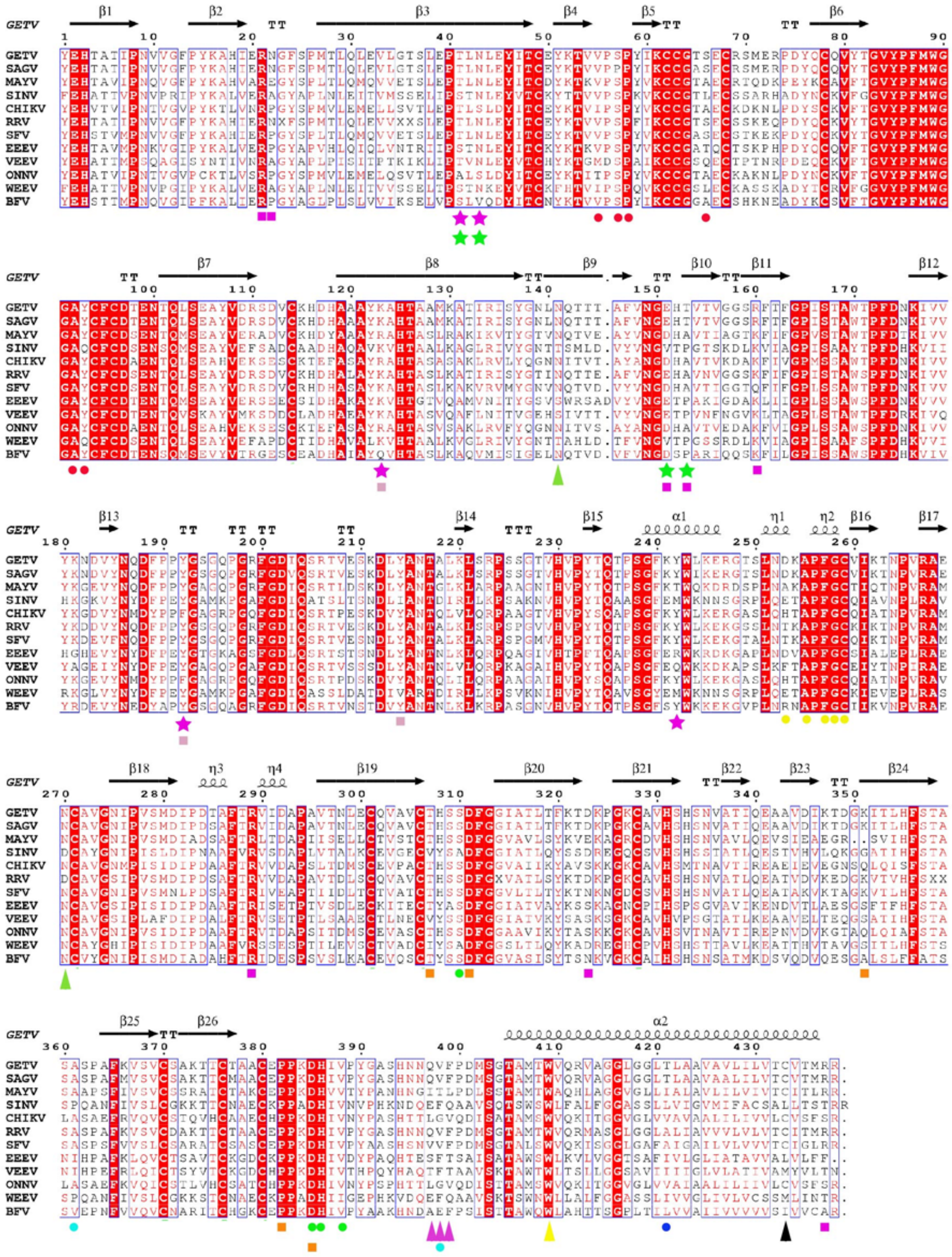

**Figure S5. Multiple sequence alignment and secondary structural elements of alphavirus E1 protein.**

Sequence alignment of E1 protein from representative members of alphavirus. The secondary structure elements are displayed above the sequences. Fully conserved residues and similar residues are shaded and shown in red. Sequences of structural polyprotein are from GETV (Getah virus, this study, ASA40294.1), SAGV (Sagiyama virus, BAA92847.1), MAYV (Mayaro virus, AZM66144.1), SINV (Sindbis virus, AYM45056.1), CHIKV (Chikungunya virus, NP_690589.2), RRV (Ross River virus, NP_062880.1), SFV (Semliki Forest virus, NP_463458.1), EEEV (Eastern equine encephalitis virus, NP_632022.1), VEEV (Venezuelan equine encephalitis virus, NP_040824.1), ONNV (O'nyong nyong virus, NP_041255.1), WEEV (Western equine encephalitis virus, NP_640331.1), and BFV (Barmah Forest virus, NP_054024.1). N-glycosylation sites and S-acylation sites are marked by green and black triangles. Magenta and red triangles represent residues that interact with cholesterol and DOPC respectively. Colored dots represent residues that interact between E1 and E2, or between E2 and capsid. Dot colors are same as the borders in Figures 2B-2E. Colored stars represent residues that interact within the ASU. Star colors are same as the corresponding residues in Figures 3B-3F. Colored squares represent residues that interact between two ASUs. Square colors are the same as the corresponding residues in Figures 4B-4G.

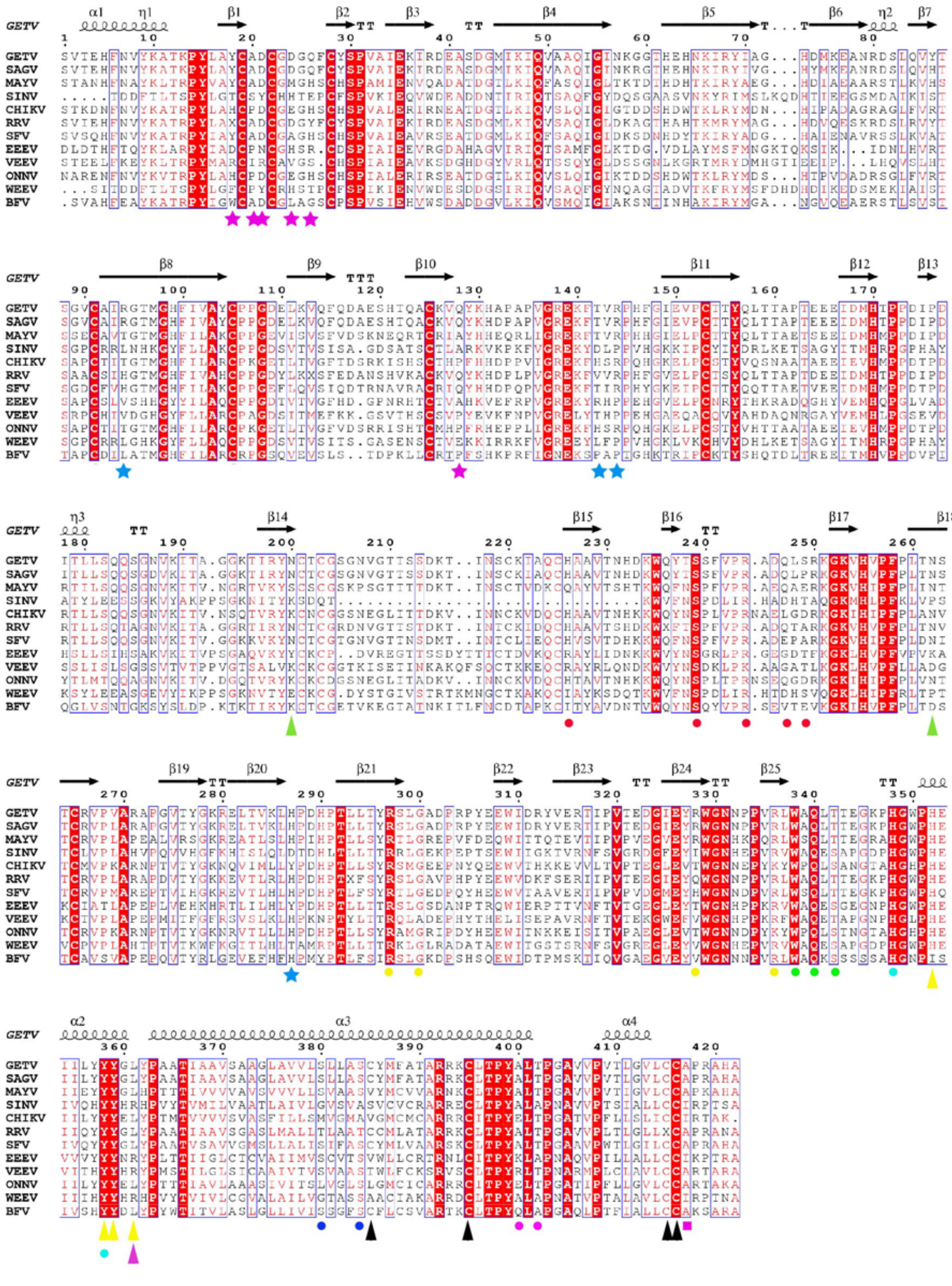

**Figure S6. Multiple sequence alignment and secondary structural elements of alphavirus E2 protein.**

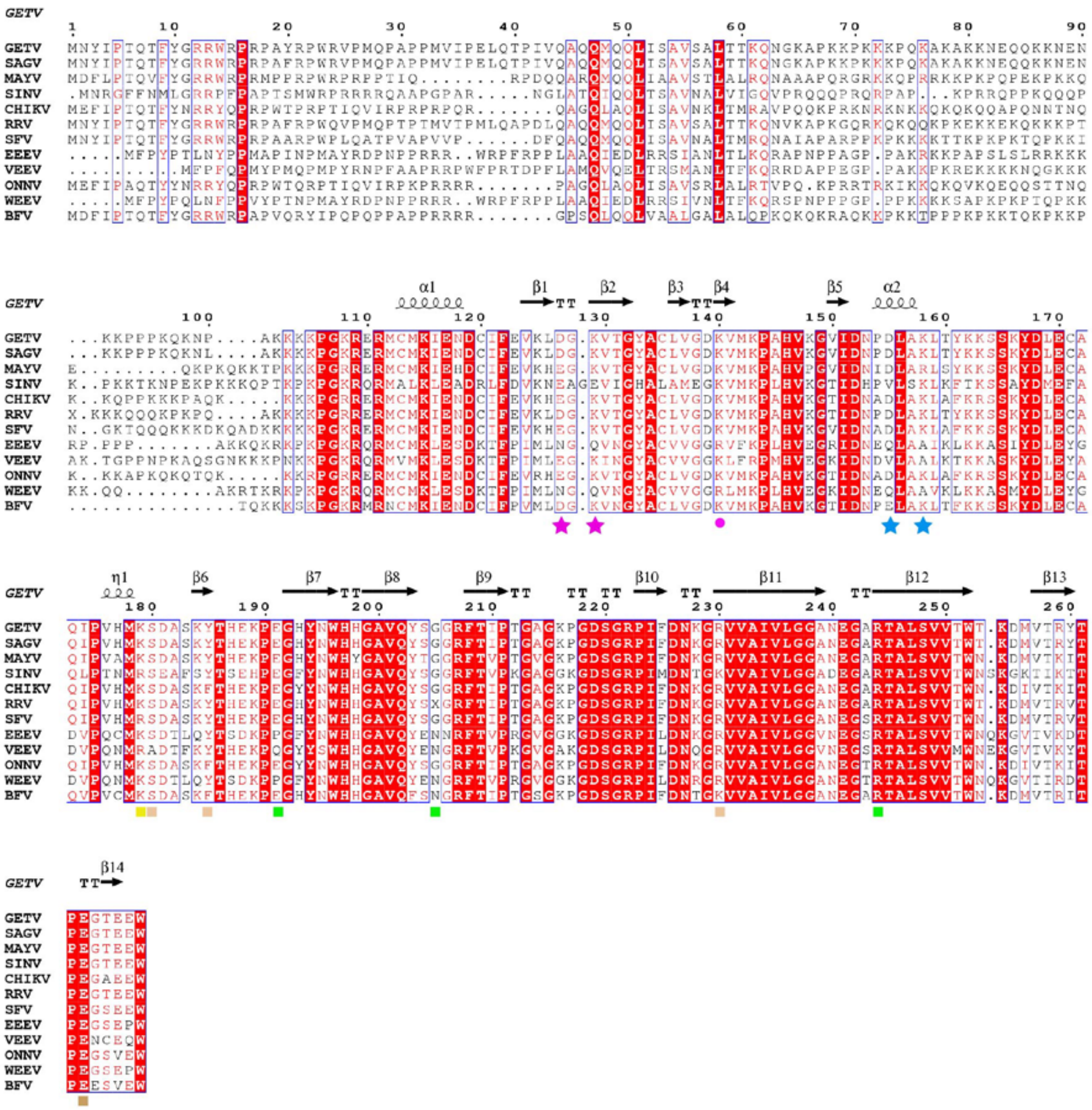

**Figure S7. Multiple sequence alignment and secondary structural elements of alphavirus capsid protein.**

**Supplemental Table 1. Cryo-EM data collection and processing, block-based reconstruction, model building, and refinement statistics.**

| **Data Collection & Processing** | | | |
|---|---|---|---|
| Microscope | FEI Titan Krios | | |
| Camera | Gatan K3 | | |
| Magnification | 105K | | |
| Voltage(kV) | 300 | | |
| Total dose ($e^{-}/Å^2$) | 40 | | |
| Defocus range (μm) | 0.8-1.5 | | |
| Pixel size (Å/pixel) | 0.83 | | |
| Symmetry imposed | I3 | | |
| Movies (total) | 16,894 | | |
| Initial particles images | 171,059 | | |
| Final particles images | 100,226 | | |
| **Block-based Reconstruction** | | | |
| Block items | Clip1(5-fold) | Clip2(3-fold) | Clip3(2-fold) |
| Block symmetry imposed | C1 | | |
| Initial block images | 6,013,560 | | |
| Final block images | 2,041,957 | 2,446,428 | 2,889,370 |
| Final resolution (Å) | 2.81 | 2.92 | 2.85 |
| **Asymmetric Unit Model Building & Refinement** | | | |
| Initial model used (PDB code) | 3J0C | | |
| Composition | | | |
| Atoms | 32,620 | | |
| Residues | 4060 amino acids | | |
| Water | 0 | | |
| Ligand | STE, PLM, PCW, CLR, NAG, BMA, MAN | | |
| Bonds (RMSD) | | | |
| Length (Å) (# > 4σ) | 0.036 | | |
| Angles (°) (# > 4σ) | 2.021 | | |
| Clash score | 2.85 | | |
| Rotamer outliers (%) | 1.44 | | |
| Ramachandran plot (%) | | | |
| Outliers | 0.30 | | |
| Allowed | 2.80 | | |
| Favored | 96.90 | | |

**Supplemental Table 2. Protein-protein interactions in GETV and other alphaviruses**

| GETV | | | | Other Alphaviruses | | | | |
|---|---|---|---|---|---|---|---|---|
| **In the E1-E2-Capsid heterotrimer** | | | | | | | | |
| **Figure** | **Interaction** | **E2** | **E1** | **Alphavirus** | **Interaction** | **E2** | **E1** | **References** |
| Fig. 2B | HB | H226 | A92 | CHIKV | HB | H226 | A92 | (Voss et al., 2010) |
| | HB | S239 | S57 | CHIKV | HB | S239 | S57 | Voss et al., 2010 |
| | HB | R244 | P58 | CHIKV | HB | R244 | P58 | Voss et al., 2010 |
| | HB | Q247 | S66 | | | | | |
| | HB | S249 | V55 | | | | | |
| Fig. 2C | HB | R297 | A255 | MAYV | HB | R297 | A255, N252 | (Ribeiro-Filho et al., 2021) |
| | HB | G300 | F257 | CHIKV | HB | G301 | F257 | Voss et al., 2010 |
| | SB | R328 | D253 | | | | | |
| | HB | R336 | G258 | CHIKV | VDM | K337 | I387, N389 | Voss et al., 2010 |
| Fig. 2D | HB | W338 | V388 | CHIKV | HB | Y339 | V388 | Voss et al., 2010 |
| | HB | Q340 | S310, D385, H386 | CHIKV | SB | Q341 | S310, D385, H386 | Voss et al., 2010 |
| | HB | T342 | H386 | | | | | |
| Fig. 2E | HB | H348 | A361 | MAYV | HB | H348 | T403 | Ribeiro-Filho et al., 2021 |
| | HB | Y358 | V398 | | | | | |
| Fig. 2F | HB | S380 | T421 | | | | | |
| | HB | S384 | T421 | | | | | |
| **Figure** | **Interaction** | **E2** | **Capsid** | **Alphavirus** | **Interaction** | **E2** | **Capsid** | **References** |
| Fig. 2G | HB | A400 | K140 | | | | | |
| | HB | T402 | K140 | | | | | |
| **In the asymmetric unit** | | | | | | | | |
| **Figure** | **Interaction** | **E2** | **E2** | **Alphavirus** | **Interaction** | **E2** | **E2** | **References** |
| Fig. 3B | HB | Y18 | R144 | CHIKV | VDW | H18 | H142, S143, R144, P145, Q146 | Voss et al., 2010 |
| | HB | A20 | R144 | CHIKV | VDW | P20 | H142, S143, R144, P145, Q146 | Voss et al., 2010 |

| | | | | | | | | |
|---|---|---|---|---|---|---|---|---|
| | SB | D21 | R144 | CHIKV | VDW | D21 | H142, S143, R144, P145, Q146 | Voss et al., 2010 |
| | SB | D24 | R94 | CHIKV | VDW | E24 | H142, S143, R144, P145, Q146 | Voss et al., 2010 |
| | HB | Q26 | R94 | CHIKV | VDW | H26 | H142, S143, R144, P145, Q146 | Voss et al., 2010 |
| | HB | Q128 | T142 | CHIKV | VDW | P128 | H142, S143, R144, P145, Q146 | Voss et al., 2010 |
| | HB | T142 | Q128 | CHIKV | VDW | H142 | H18, P20, D21, E24, G25, H26, S27, E109, T110, T126, P128, | Voss et al., 2010 |
| **Figure** | **Interaction** | **E2** | **E1** | **Alphavirus** | **Interaction** | **E2** | **E1** | **References** |
| Fig. 3C | HB | H287 | Y242 | CHIKV | VDW | Y288 | Q218, S234, Q235, A236, P237, | Voss et al., 2010 |
| **Figure** | **Interaction** | **E1** | **E1** | **Alphavirus** | **Interaction** | **E1** | **E1** | **References** |
| Fig. 3D | SB | K123 | E151 | CHIKV | VDW | R123 | N149, D151, H152 | Voss et al., 2010 |
| | HB | Y192 | E151, T153 | CHIKV | VDW | F192 | Y147 | Voss et al., 2010 |
| Fig. 3E | HB | T41 | N43 | SFV | HB | T41 | N43 | (Roussel et al., 2006) |
| | HB | N43’ | T41’ | SFV | HB | N43 | T41 | Roussel et al., 2006 |
| **Figure** | **Interaction** | **Capsid** | **Capsid** | **Alphavirus** | **Interaction** | **Capsid** | **Capsid** | **References** |
| Fig. 3F | HB | D120 | K151 | | | | | |

| | | | | | | | | |
|---|---|---|---|---|---|---|---|---|
| | HB | K122 | D148 | | | | | |
| **Between two asymmetric units** | | | | | | | | |
| **Figure** | **Interaction** | **E1** | **E1** | **Alphavirus** | **Interaction** | **E1** | **E1** | **References** |
| Fig. 4B | HB, SB | E151 | K123, Y192 | CHIKV | VDW | D151 | R123, K176, P191, W192, G193, A194, Y214, R206 | Voss et al., 2010 |
| | HB | T153 | Y192 | CHIKV | VDW | A153 | W192, G193, A194 | Voss et al., 2010 |
| | HB | R160 | Y192 | | | | | |
| | HB | R160 | Y214 | | | | | |
| Fig. 4C | SB | R21 | D385 | SFV | SB | R21 | D284 | Roussel et al., 2006 |
| | HB | N22 | T307, P382 | CHIKV | VDW | P22 | T307, H381, P382, P383, R384 | Voss et al., 2010 |
| | SB | R289 | D311 | | | | | |
| Fig. 4D | SB | D323 | K351 | CHIKV | VDW | S323 | V315, I317, K319, E353 | Voss et al., 2010 |
| **Figure** | **Interaction** | **Capsid** | **Capsid** | **Alphavirus** | **Interaction** | **Capsid** | **Capsid** | **References** |
| Fig. 4E | HB | S180 | R244 | | | | | |
| | HB | Y185 | G205 | | | | | |
| | SB | R230 | E191 | | | | | |
| Fig. 4F | SB | E263 | K179 | | | | | |
| **Figure** | **Interaction** | **E2** | **E1** | **Alphavirus** | **Interaction** | **E2** | **E1** | **References** |
| Fig. 4G | HB | A417 | R437 | | | | | |

HB, Hydrogen bonds; SB, salt bridges; VDM, van der Waals contact.
Contacts were calculated using PISA (Lawrence and Colman, 1993).
Interactions highlighted in red represent the newly discovered interactions in the present study.

**Supplemental Table 3. Glycosylation sites in E1 and E2 proteins**

| Protein | Glycosylation Type | Method | Glycosylation Site |
|---|---|---|---|
| E1 | O-Glycosylation | NetOGlyc (Steentoft et al., 2013) (www.cbs.dtu.dk/services/NetOGlyc/) | S250 |
| | | YinOYang (Gupta and Brunak, 2002) (www.cbs.dtu.dk/services/YinOYang/) | T53, **S66**, S104, T132, T144, S222, S226, T234, T288, S297, T368, S377, T435 |
| | | Previously reported | SINV S238 (no density) (Ribeiro-Filho et al., 2021) |
| | | **Cryo-EM density (no atomic model)** | **S66** |
| | N-Glycosylation | NetNGlyc (Gupta and Brunak, 2002) (www.cbs.dtu.dk/services/NetNGlyc/) | N9, N22, N43, N100, N139, **N141,** N149, N175, N182, N186, N216, N252, N264, **N270**, N275, N298, **N335**, N395, N396. |
| | | GlycoMine (Li et al., 2015) (glycomine.erc.monash.edu/Lab/GlycoMine/) | N9, N22, N43, N100, N139, **N141,** N149, N175, N182, N186, N216, N252, N264, **N270**, N275, N298, N395, N396. |
| | | **Mass spectrometry** | N9, N100, N139, **N141**, N149, N252, N264, **N270**, N275, N395, N396. |
| | | Previously reported | SINV N139, N245 (density, no atomic model) (Chen et al., 2018)<br>**CHIKV, MAYV N141 (atomic model)** (Ribeiro-Filho et al., 2021; Voss et al., 2010) |
| | | **Cryo-EM density (no atomic model)** | **N335** |
| | | **Cryo-EM density (atomic model)** | **N141, N270** |
| E2 | O-Glycosylation | NetOGlyc | T142, T154, **T155**, T159, S160, S163. |
| | | YinOYang | T3, T87, S88, T122, S185, T211, S212, S213, T230, T238, S240, T261, **T264**, T343, T366. |
| | | Previously reported | N/A |
| | | **Cryo-EM density (no atomic model)** | **T155, T264** |

| | | | |
|---|---|---|---|
| | N-Glycosylation | NetNGlyc | N7, N33, N57, N65, N79, N187, **N200**, N207, N218, N231, **N262**, N332. |
| | | GlycoMine | N7, N57, N65, N79, N187, **N200**, N207, N218, N231, **N262**, N331, N332. |
| | | **Mass spectrometry** | N57, N187, **N200**, N218, N231, **N262**. |
| | | Previously reported | **CHIKV N196 (atomic model)** (Voss et al., 2010)<br>**MAYV N262 (atomic model)** (Ribeiro-Filho et al., 2021)<br>SINV N283 (density, no atomic model) (Chen et al., 2018) |
| | | **Cryo-EM density (atomic model)** | **N200, N262** |

**Supplemental Table 4. S-acylation sites in E1 and E2 proteins**

| Protein | Acylation Type | Method | S-acylation Site |
|---|---|---|---|
| **E1** | S-acylation | CSS-Palm (Ren et al., 2008) (www.csspalm.biocuckoo.org/online.php) | C62, C63 |
| | | GPS-Palm (Ning et al., 2021) (gpspalm.biocuckoo.cn/) | C63, C259, **C433** |
| | | Mass spectrometry | C62, C63, C68, C78, C271, **C433** |
| | | Previously reported | BFV C436 (density, no atomic model) (Kostyuchenko et al., 2011) |
| | | **Cryo-EM density (atomic model)** | **C433** |
| | N-acylation, O-acylation | Mass spectrometry | S136, T143 |
| | | Previously reported | N/A |
| | | **Cryo-EM density** | **N/A** |
| **E2** | S-acylation | CSS-Palm | C19, C22, **C415**, **C416** |
| | | GPS-Palm | **C385**, **C395**, **C415**, **C416** |
| | | Mass spectrometry | C22, C91, C105, C153, C203, C225, **C385**, **C395**, **C415**, **C416** |
| | | Previously reported | BFV C384, C387, C394, C414, C415 (density, no atomic model) (Kostyuchenko et al., 2011)<br>VEEV C396, C416, C417 (density, no atomic model) (Zhang et al., 2011) |
| | | **Cryo-EM density (atomic model)** | **C385, C395, C415, C416** |
| | N-acylation, O-acylation | Mass spectrometry | T130, T159, T171, K195, T196 |
| | | Previously reported | N/A |
| | | **Cryo-EM density** | **N/A** |
