## Supplementary material for "Structure of Infective Getah Virus at 2.8 Å-resolution Determined by Cryo-EM": Star Methods

Detailed methods are provided in the online version of this paper and include the following:

- KEY RESOURCES TABLE
- RESOURCE AVAILABILITY
  - Lead Contact
  - Materials Availability
  - Data Availability
- METHOD DETAILS
  - Virus infection
  - Virus production and purification
  - Cryo-EM sample preparation and data acquisition
  - Image processing
  - Model building and refinement
  - Quantitative N-glycosylation analysis by LC-MS/MS
  - Quantitative S-acylation analysis by LC-MS/MS
  - Molecular dynamics simulation study of cholesterols and DOPC regulation of E1-E2

### KEY RESOURCES TABLE

| REAGENT or RESOURCE | SOURCE | IDENTIFIER |
|---|---|---|
| **Antibodies** | | |
| K1G4F4 | This paper | Patent ID: CN201910649881.2 |
| **Bacterial and Virus Strains** | | |
| **GETV-V1** | http://pubmed.ncbi.nlm.nih.gov/32781023/ | GenBank: KY399029.1 |
| **Chemicals, Peptides, and Recombinant Proteins** | | |
| Trypsin | Promega | Cat#5280 |
| rPNGase F | New England Biolabs | Cat#P0709 |
| **Critical Commercial Assays** | | |
| GETV RT-PCR | Zhou et al., 2020 | http://pubmed.ncbi.nlm.nih.gov/32781023/ |
| **Deposited Data** | | |
| CryoEM map of native GETV (2.8 Å) | This paper | EMD-31533 |
| Cryo-EM map of the clipped virus at the 5-fold axis of asymmetry (2.81 Å) | This paper | EMD-31614 |

| | | |
|---|---|---|
| Cryo-EM map of the clipped virus at the 3-fold axis of asymmetry (2.92 Å) | This paper | EMD-31615 |
| Cryo-EM map of the clipped virus at the 2-fold axis of asymmetry (2.85 Å) | This paper | EMD-31617 |
| Coordinates of E1-E2-Capsid protein for asymmetric unit | This paper | PDB ID 7FD2 |
| **Experimental Models: Cell Lines** | | |
| BHK21 | ATCC | Cat#CCL-10 |
| **Experimental Models: Organisms/Strains** | | |
| Mouse：BALB/c | The Jackson Laboratory | JAX: 000651 |
| **Software and Algorithms** | | |
| Prism | GraphPad | http://www.graphpad.com/ |
| NetOGlyc 4.0 | Steentoft et al., 2013 | http://www.cbs.dtu.dk/services/NetOGlyc/ |
| YinOYang 1.2 | Gupta et al., 2002 | http://www.cbs.dtu.dk/services/YinOYang/ |
| NetNGlyc 1.0 | Gupta et al., 2002 | http://www.cbs.dtu.dk/services/NetNGlyc/ |
| GlycoMine | Li et al., 2015 | http://glycomine.erc.monash.edu/Lab/GlycoMine |
| CSS-Palm | Ren et al., 2008 | http://www.csspalm.biocuckoo.org/online.php |
| GPS-Palm | Ning et al., 2021 | http://gpspalm.biocuckoo.cn/ |
| Mitioncor2 | Zheng et al., 2017 | http://emcore.ucsf.edu/ucsf-software |
| CTFFIND4 | Rohou and Grigorieff et al., 2017 | http://grigoriefflab.umassmed.edu/ctffind4 |
| Relion | Bharat and Scheres, 2016 | http://scicomp.ethz.ch/wiki/RELION |
| UCSF Chimera | Pettersen et al., 2004 | http://www.rbvi.ucsf.edu/chimera/ |
| jspr | Guo and Jiang et al., 2004 | http://jiang.bio.purdue.edu/jspr/ |
| Phenix Real Space Refine | Adams et al., 2010 | http://strucbio.biologie.uni-konstanz.de/ccp4wiki/index.php?title=Phenix |
| MolProbity | Chen et al., 2010 | http://molprobity.manchester.ac.uk/ |
| COOT | Emsley and Cowtan,2004 | http://strucbio.biologie.uni-konstanz.de/ccp4wiki/index.php?title=Coot |
| PyMOL | Schrödinger, LLC | http://pymol.org/2/ |
| ExPasy | Gasteiger et al., 2003 | http://www.expasy.org |
| VMD | Humphrey et al., 1996 | http://www.ks.uiuc.edu/Research/vmd/ |
| CHARMM-GUI | Jo et al., 2008 | http://www.swmath.org/software/21732 |
| H++ | Gordon et al, 2005 | http://biophysics.cs.vt.edu/ |

| | | |
|---|---|---|
| AMBER 2019 | Case et al., 2019 | http://ambermd.org/ |
| gunplot 5.2 | Merritt et al., 2019 | http://sourceforge.net/projects/gnuplot/ |
| **Other** | | |
| 300-mesh, R2/1, amorphous alloy film | CryoMatrix | Cat#M025-Au300-R20/10 |

### RESOURCE AVAILABILITY

#### ○ Lead Contact

#### ○ Materials Availability

Viral strain and antibody are available from Chuanqing Wang with a completed Materials and Transfer Agreement.

#### ○ Data Availability

The atomic coordinates of E1-E2-Capsid of the GETV have been deposited in the Protein Data Bank under accession code 7FD2. The cryo-EM map of the envelope glycoprotein and capsid core, and the clipped virus at 5-fold, 3-fold and 2-fold have been deposited in the Electron Microscopy Data Bank under accession code EMD-31533, EMD-31614, EMD-31615, and EMD-31617.

### METHOD DETAILS

#### ○ Virus infection

All protocols were approved by the Institutional Animal Care and Use Committee of Shanghai Tenth People's Hospital. Mice were apparently healthy and had no serum neutralizing antibody against GETV before the experiment. Newborn mice (2-day old or 3-day old) were inoculated oronasally with 10μl ($10^6$ $TCID_{50}$/mL, 50% tissue culture infectious dose) GETV-V1 strain. Pregnant mice were inoculated oronasally with 100 μl ($10^6$ $TCID_{50}$/mL) GETV-V1 strain in early-gestation (embryonic day 6, E6), middle-gestation (E10), and late-gestation (E14).

#### ○ Virus production and purification

Mature and infective GETV virions were handled in BSL2 facilities at Cryo-Electron Microscopy Center, Southern University of Science and Technology, and at College of Animal Veterinary Medicine, Henan Agricultural University. GETV-V1 strain (GenBank Sequence Accession: KY399029.1) was incubated in monolayers Baby Hamster Kidney

Fibroblast Cells (BHK-21, ATCC CCL-10) at an approximate multiplicity of infection (MOI) of 0.1 for 30 min to allow viruses to bind and enter into the BHK cells. The supernatant was removed and replaced with fresh culture medium (DMEM with 2% FBS and 1% PS). GETV were proliferated in 10 T175 culture flasks and harvested 36 h post infection. The cell debris of the supernatant was removed at 10,000 × g for 1h (Sorvall LYNX 6000 Superspeed Centrifuge). The GETV particles were concentrated by centrifuging at 80,000 × g for 1.5 h in Ultracentrifuge (Beckman Optima XPN-100) with a type 45 Ti rotor at 4 °C. After soaking in 30 ml phosphate-buffered saline (PBS, pH 7.47) for 4h, the virus suspension was initially purified by through a 20% (w/v) sucrose cushion at 80,000 × g for 1.5 h in a type SW32 Ti rotor at 4 °C. The virus pellet was gently resuspended in 2 ml PBS buffer for 4h and loaded onto a linear 20-50% (w/v) sucrose density gradient at 100,000 × g for 2 h in a SW41 Ti rotor at 4 °C. The light scattering band corresponding to the virus particles collected and resuspended in PBS. The integrity and purity of the GETV particles were examined by SDS-PAGE and mass spectrometry, and by negative-stain electron microscopy.

#### ○ Cryo-EM sample preparation and data acquisition

A 5 μl virus sample was added to the glow-discharged 400-mesh grid covered with amorphous nickel-titanium alloy film (ANTA film, R2/1) at 100% humidity and 6 °C, blotted with filter paper and frozen by plunging into liquid ethane using a Vitrobot Mark IV system (Thermo Fisher Scientific Inc.). Cryo-EM data were collected on a 300 kV Titan Krios microscope (Thermo Fisher Scientific Inc.) equipped with a BioContinuum Imaging Filter (Gatan Inc). The images were recorded on a K3 summit direct detection camera (Gatan Inc) using SerilEM software for automated image acquisition. The images were recorded at a nominal magnification of 105,000 × in super-resolution mode, yielding a calibrated pixel size of 0.83 Å. Each exposure was dose-fractionated into 32 frames leading to a total dose of 40 $e^-/Å^2$. The final defocus range of the micrographs was -0.8 to -1.5 μm.

#### ○ Image processing

A total of 16,894 movie stacks of the GETV were collected. The beam-induced drift was corrected using MotionCor2 (Zheng et al., 2017). The contrast transfer function (CTF) parameters were estimated using CTFFIND4 (Rohou and Grigorieff, 2015). In total, 171,059 particles were extracted using the EMAN2 (Tang et al., 2007). The 4× binned particle images were subjected to reference-free 2D classification in RELION-3 (Zivanov et al., 2018). After several rounds of reference-free 2D classification, a subset of 144,254 particles was isolated for 3D classification and reconstruction. Two maps at 8.94 Å-resolution and 9.54 Å-resolution were obtained with 56,573 and 43,653 good particles, respectively. The coordinates were used to extract the 2× binned particles for 3D refinement using RELION-3 with I3 symmetry imposed, which resulted in two maps at 6.64 Å-resolution and 6.69 Å-resolution obtained with 43,775 and 62,415 good particles, respectively. The rotational and translational parameters of each particle were further refined with jalign program from JSPR, which then enabled the reconstruction of a better map at 4.1 Å-resolution using the j3dr program in JSPR (Guo and Jiang, 2014). To overcome the defocus gradient and the heterogeneity of the ~70nm virus, block-based reconstruction was performed to further improve the resolution (Zhu et al., 2018). Three types of block (five trimers near icosahedral five-fold axis, four trimers near icosahedral-three-fold axis and two trimers near icosahedral two-fold axis) were selected, 3D classified and refined separately. Maps at 3.05 Å, 3.28 Å, and 3.14 Å, were obtained using unbinned particles. At this stage, CTF refinement was carried out to better estimate the local defocus value for each block, and a final round of 3D refinement generated density maps in resolution of 2.81 Å, 2.92 Å, and 2.85 Å, respectively, as estimated by the gold-standard Fourier shell correlation (FSC) cut-off value of 0.143 (**Figure S3**). The local resolution distribution of the final reconstruction was assessed using ResMap (Kucukelbir et al., 2014) (**Figure S3**).

#### ○ Model building and refinement

An asymmetric subunit of VEEV model (PDB ID: 3J0C) was used as template and rigid-body fitted into the 2.8 Å GETV map using UCSF Chimera (Pettersen et al., 2004). The amino acid sequences of VEEV were then mutated to GETV and an initial 3D atomic model of E1–E2-Capsid was built using Coot (Emsley et al., 2010). The resulting GETV coordinates were then used as a starting point to a flexible refinement through the PHENIX (Adams et al., 2010). The refinement cycle was repeated, and the quality of the final 3D atomic model of E1-E2-Capsid was evaluated using MolProbity (Chen et al., 2010). The Cryo-EM data collection and processing, block-based reconstruction, model building, and refinement statistics are summarized in **Table S1**.

#### ○ Quantitative N-glycosylation analysis by LC-MS/MS

E1 and E2 gel bands were excised for LC-MS/MS for quantitative N-glycosylation analysis. Protein bands were dissolved by 200μl of 8M urea with 10mM DTT and digested by 100μl of 25mM ammonium bicarbonate containing 0.01μg/μl trypsin before being collected and lyophilized in a sterile centrifuge. The dried polypeptide was dissolved in 40mM $NH_4HCO_3$ prepared with O18. Two units of rPNGase F was defined as the amount of enzyme that digests 100μg polypeptide. The lyophilized peptide fractions were re-suspended in dd$H_2O$ containing 0.1% formic acid. The online Chromatography separation was performed on the EASY-nLC1200 system. The trapping and desalting procedure were carried out with 20μl solvent A (0.1% formic acid). DDA mass spectrum techniques were used to acquire tandem MS data on a Thermo Fisher™ Q Exactive™ Mass Spectrometer fitted with a Nano Flex ion source. For a full mass spectrometry survey scan, the target value was $3\times10^6$ and the scan ranged from 350 to 2,000m/z at a resolution of 70,000. For the MS2 scan, only spectra with a charge state of 2-5 were selected for fragmentation by higher-energy collision dissociation. N-linked glycans can be released from the glycoprotein with the enzyme N-glycosidase F rPNGase F. Asparagine residues can cause a mass increase of 2.9882 Da in “heavy” water and identified the N-glycosylation sites by LC-MS/MS peptide mapping. The MS data were analyzed for protein modification using PEAKS Studio 8.5.

#### ○ Quantitative S-acylation analysis by LC-MS/MS

E1 and E2 gel bands were excised for LC-MS/MS for quantitative S-acylation analysis.100μl of decolorizing solution was added and placed at room temperature to decolor overnight. 100% acetonitrile was added until the gel mass turned white. 10μl 0.01μg/μl trypsin was added to the gel to fully absorb by iced until it became transparent. The concentrated and dried trypsin-cut peptide samples were re-dissolved in ddH2O containing 0.1% formic acid and analyzed by on-line LC-MS/MS. The online chromatography separation was performed on the EASY-nLC1200 system. The trapping and desalting procedure was carried out with 20μl solvent A (0.1% formic acid). DDA mass spectrum techniques were used to acquire tandem MS data on a Thermo Fisher Fusion Lumos Mass Spectrometer fitted with a Nano Flex ion source. For a full mass spectrometry survey scan, the target value was $3\times10^6$ and the scan ranged from 300 to 1,800m/z at a resolution of 70,000. For the MS2 scan, only spectra with a charge state of 2-7 were selected for fragmentation by higher-energy collision dissociation. S-acylation modification occurs on CKST amino acids, for which the molecular weight change after modification is 238.23/266.26 Da (palmitic acid/stearic acid). The MS data were analyzed for protein modification using Proteome Discoverer 2.4.

#### ○ Molecular dynamics simulation study of cholesterols and DOPC regulation of E1-E2

The solved cryo-EM structure of the full length E1-E2 complex was first truncated first to keep only the amino acids of 396-438 for E1 and 340-403 for E2 as the starting protein model for MD simulations. The truncated protein (denoted as E1-E2), together with the ligands at three different compositions, was then packed with the lipid bilayer using the CHARMM-GUI tool (Lee et al., 2019). The resulting three systems to simulate are E1-E2 with: (1) all the cholesterols and DOPC molecule were kept (denoted as Wild Type); (2) removal of the pocket cholesterol only (denoted as delCHL); (3) removal of the pocket DOPC only (denoted as delDOPC); (4) removal of both cholesterol and DOPC in the pocket (denoted as delCHL/DOPC); (5) removal of the two cholesterols surround the pocket (denoted as del2CHL); and (6) all 3 cholesterols and DOPC molecule removed (denoted as delALL). The main lipid components with the same mixed lipid ratio in both inner and outer leaflets (DPPC/DOPC/POPC/DPPE/POPE/DPPS/POPS/Chol=7.5:7.5:15:5:7:5:7:46) were used in constructing the membrane model following the reported lipidomic data (Kalvodova et al., 2009a). Here the sphingomyelin was approximatively modeled by the POPC due to the lack of the force filed parameters for sphingomyelin and the identical hydrophilic phosphorylcholine head group, similar conformation and charge distribution between the two molecules. In order to obtain a consistent protein-membrane packing conformation with the cryo-EM density data, the vector from the residue Val-410 of E1 to Lys-394 of E2 was aligned with the normal axis of the lipid bilayer when packing systems. Next, the constructed protein-ligand-membrane systems were submitted to the H++ web server (Gordon et al., 2005) to determine the protonation states of the protein under the membrane environment in order to approximate a simulation pH of 7.0. As a result, the amino acids His-355 in E1 and His-352 in E2 were simulated as pronated. The protein was diagonally positioned in the X-Y plane in the simulation box filled with the TIP3P water model (Jorgensen et al., 1983) to save space, leading to a box size of ~87 Å × 87 Å × 122 Å. The final systems were neutralized with the sodium ion and extra NaCl was added to model the experimental salt concentration of 0.15 M. All MD runs were performed using AMBER 19 package (Case et al., 2005) with the ff14SB protein force field (Maier et al., 2015) and the LIPID17 lipid force field (Dickson et al., 2014). The van de Waals cutoff was set to 10 Å and PME was used for long-range electrostatic energy calculation. Minimization, heating and holding processes were carried out before production run. All production simulations were performed at 310 K under the NPT ensemble with the integration step of 2 fs and an anisotropic pressure coupling. Five independent runs for each system of Wild Type, delCHL, delDOPC, delCHL/DOPC, del2CHL, or delALL were carried out to improve samplings, which led to a total of 30 simulations. Production trajectories of 700 ns were generated for all simulations. All figures and animations were prepared using gnuplot 5.4 and VMD (Humphrey et al., 1996).
